# Brassicaceae display diverse photorespiratory carbon recapturing mechanisms

**DOI:** 10.1101/2022.12.22.521581

**Authors:** Urte Schlüter, Jacques W. Bouvier, Ricardo Guerreiro, Milena Malisic, Carina Kontny, Philipp Westhoff, Benjamin Stich, Andreas P. M. Weber

**Affiliations:** Institute of Plant Biochemistry, Cluster of Excellence for Plant Sciences (CEPLAS), Heinrich Heine University, Universitätsstr. 1, 40225 Düsseldorf, Germany; Institute for Quantitative Genetics and Genomics of Plants, Cluster of Excellence for Plant Sciences (CEPLAS), Heinrich Heine University, Universitätsstr. 1, 40225 Düsseldorf, Germany; Metabolomics and Metabolism Laboratory, Cluster of Excellence for Plant Sciences (CEPLAS), Heinrich Heine University, Universitätsstr. 1, 40225 Düsseldorf, Germany; Department of Plant Sciences, University of Oxford, South Parks Road, Oxford OX1 3RB, UK; Department of Plant Microbe Interactions, Max Planck Institute for Plant Breeding Research, 50829 Cologne, Germany

**Keywords:** carbon concentrating mechanisms, photorespiration, C_3_-C_4_ intermediacy, C_2_ photosynthesis, photorespiratory glycine shuttle, Brassicaceae, CO_2_ compensation point (CCP), bundle sheath (BS), mesophyll (M)

## Abstract

Carbon concentrating mechanisms enhance the carboxylase efficiency of the central photosynthetic enzyme rubisco by providing supra-atmospheric concentrations of CO_2_ in its surrounding. In the C_4_ photosynthesis pathway, this is achieved by combinatory changes to leaf biochemistry and anatomy. Carbon concentration by the photorespiratory glycine shuttle requires fewer and less complex modifications. It could represent an early step during evolution from C_3_ to C_4_ photosynthesis and an inspiration for engineering approaches. Plants displaying CO_2_ compensation points between 10 to 40 ppm are therefore often termed ‘C_3_–C_4_ intermediates’. In the present study, we perform a physiological, biochemical and anatomical survey of a large number of Brassicaceae species to better understand the C_3_-C_4_ intermediate phenotype. Our phylogenetic analysis suggested that C_3_-C_4_ metabolism evolved up to five times independently in the Brassicaceae. The efficiency of the pathways showed considerable variation between the species but also within species. Centripetal accumulation of organelles in the bundle sheath was consistently observed in all C_3_-C_4_ classified accessions indicating a crucial role of anatomical features for CO_2_ concentrating pathways. Leaf metabolite patterns were strongly influenced by the individual plant accessions, but accumulation of photorespiratory shuttle metabolites glycine and serine was generally observed. Analysis of PEPC activities suggests that C_4_-like shuttles have not evolve in the investigated Brassicaceae.

**Highlight:** Our physiological, biochemical and anatomical survey of Brassicaceae revels multiple evolution of C_3_-C_4_ intermediacy connected to variation in photorespiratory carbon recapturing efficiency and a distinct C_3_-C_4_ bundle sheath anatomy.

## Introduction

The majority of plant species on Earth, including many crops, employ C_3_ photosynthesis. In these plants, under the present environmental conditions, the central photosynthetic enzyme rubisco (ribulose-1,5-bisphosphatate carboxylase/oxygenase) fixes approximately one molecule of oxygen for every three molecules of CO_2_ (Sharkey, 1988). Here, whilst the carboxylase reaction of rubisco produces two molecules of 3-phosphglycerate (3PGA) which feeds into the Calvin-Benson-Bassham cycle (CBB), the oxygenase reaction produces 2-phosphoglycolate (2PG). 2PG is a competitive inhibitor of some CBB enzymes (Flügel *et al*., 2017) and hence must be rapidly removed. Further, carbon contained in 2PG must be recycled into 3PGA to prevent depletion of CBB intermediates. These functions are fulfilled by the photorespiratory pathway. Photorespiration consists of coordinated enzyme activities that are located in different cellular compartments. In the plastids, 2PG is converted into glycolate followed by oxidation and transamination in the peroxisome producing glycine. Glycine is transported into the mitochondria and metabolised into serine by glycine decarboxylation. Serine is finally converted into glycerate in the peroxisome and into 3PGA in the plastid. During glycine decarboxylation, previously fixed carbon and nitrogen is converted into CO_2_ and NH_3_. The re-fixation of carbon and nitrogen into organic forms, however, requires energy. Therefore, photorespiration is often considered a wasteful process in terms of energy-, carbon- and nitrogen-balance. Further, the affinity of rubisco for O_2_ increases with rising temperatures, in addition stomata closure during water shortages can lead to a drop in CO_2_ concentrations inside the leaf. Thus, climate change could contribute to increased 2PG production.

Possibilities to reduce photorespiratory losses that are being explored today include increasing the capacity of the plant to recapture photorespiratory CO_2_, modifying rubisco kinetic properties and introducing carbon concentrating mechanisms to limit the oxygenase reaction of rubisco by creating an CO_2_ rich environment around the enzyme (Busch *et al*., 2013). Better understanding of naturally occurring carbon concentration mechanisms will help in the design of biotechnological approaches.

In C_3_ species, photosynthesis and photorespiration mainly take place in the mesophyll (M) of the leaves. Recapture of photorespiratory CO_2_ can be facilitated by arrangement of a continuous layer of plastids at the cell periphery next to the intercellular space. This arrangement acts to block diffusion of CO_2_ out of the cell because any CO_2_ which would otherwise escape would now need to pass through the plastids where it can be reassimilated by rubisco (Sage and Sage, 2009; Busch *et al*., 2013). In rice, rubisco containing extensions (stromules) can increase the area of the plastidial barriers, preventing the efflux of CO_2_ produced in the mitochondria (Sage and Sage, 2009). C_3_ species like wheat and rice achieve photorespiratory reassimilation rates of 24-38% (Busch *et al*., 2013).

The number of chloroplasts in the C_3_ bundle sheath (BS) varies between species (Leegood, 2008), but they tend to be smaller and fewer in number compared to the M. Contribution of BS chloroplasts to leaf photosynthesis is considered to be small (Kinsman and Pyke, 1998; Janacek *et al*., 2009; Aubry *et al*., 2014). Nevertheless, BS cells possess important roles in leaf hydraulics, phloem loading, and intra-leaf signalling (Leegood, 2008; Lundgren *et al*., 2014). In *Arabidopsis*, a cell specific translatome analysis indicated distinct roles for the BS cells in sulphur metabolism and transport, as well as in glucosinolate and trehalose metabolism (Aubry *et al*., 2014). Enlargement of BS cells has initially been associated with improved leaf hydraulics under water limiting conditions (Sage, 2004). Associated increases in BS organelle numbers indicate enhanced photosynthetic and photorespiratory activity. Centripetal arrangement of BS organelles helps to reduce loss of photorespiratory CO_2_ (Muhaidat *et al*., 2011; Sage *et al*., 2014). Such BS cells have also been labelled as “activated” or “proto-Kranz” anatomy (Gowik and Westhoff, 2011; Sage *et al*., 2014).

Further increases in BS CO_2_ concentration are possible by shifting the glycine decarboxylation step from the M to the BS (Monson *et al*., 1984; Rawsthorne *et al*., 1988*a*; Sage *et al*., 2014). This rearrangement forces photorespiratory glycine produced in the M to diffuse to the BS, where the tissue-specific increase in glycine decarboxylation activity promotes an elevated concentration of CO_2_ around the BS rubisco and, thus, supresses its oxygenase reaction. The glycine decarboxylation step is realised by four proteins (GLDP, GLDH, GLDL, GLDT) which combined, form the glycine decarboxylase complex (GDC) that operates in coordination with the serine hydroxymethyltranserase (SHMT) (Douce *et al*., 2001). BS localised activity of glycine decarboxylation is mainly associated with cell specificity of the GLDP-protein (Rawsthorne *et al*., 1988*a*; Schulze *et al*., 2013). The installation of a glycine shuttle is accompanied by a further increase in organelle numbers in the BS. The majority of mitochondria are therefore located between the centripetally arranged plastids and the vein orientated cell wall (Sage *et al*., 2014). Such combination of carbon supply by the glycine shuttle and efficient CO_2_ scavenging by adequate organelle arrangement improves the leaf carbon conservation and can be measured as reduction in the carbon compensation point (CCP or □). Plants employing the photorespiratory glycine shuttle are often classified as C_2_ species or C_3_-C_4_ intermediates of type I (Edwards and Ku, 1987; Sage *et al*., 2014). Their CO_2_ reassimilation capacity was estimated to be around 73 % in *M. arvensis* (Hunt *et al*., 1987).

GDC activity is thought to be absent or considerably reduced in the M cells of well-developed C_2_ species, perhaps as a result of a loss-of-function mutation or insertion of a transposable element early in C_2_ evolution (Rawsthorne, 1992; Sage *et al*., 2012; Adwy *et al*., 2015). Consistently, preferential BS localisation of GLDP has been observed in many well-developed C_2_ species, including, in the dicots *Moricandia* (Rawsthorne *et al*., 1988*a*), *Flaveria* (Hylton *et al*., 1988), *Diplotaxis* (Ueno *et al*., 2003), *Cleome* (Marshall *et al*., 2007; Voznesenskaya *et al*., 2007), *Salsola* (Voznesenskaya *et al*., 2001; Wen and Zhang, 2015; Schüssler *et al*., 2017), *Brassica* (Ueno, 2011), *Euphorbia* (Sage *et al*., 2011*c*), *Heliotropium* (Muhaidat *et al*., 2011), *Anticharis* (Khoshravesh *et al*., 2012) and *Blepharis* (Fisher *et al*., 2015), as well as the monocots *Neurachne* (Christin *et al*., 2012), *Alloteropsis* (Lundgren *et al*., 2016) and *Homolepis* (Khoshravesh *et al*., 2016).

Assimilatory power of the BS can generally be further enhanced by decarboxylation of additional metabolites. Transcripts of genes encoding decarboxylating enzymes for the C_4_ metabolites aspartate and malate already preferentially accumulate in the C_3_ BS (Aubry *et al*., 2014). Organelle accumulation and enhancement of organellar metabolism in the BS could increase the availability of such compounds. The glycine shuttle transports not only carbon between M and BS, but also creates a nitrogen imbalance between these cells. For instance, two glycine molecules are converted into one serine molecule while CO_2_, NH_3_ and NADH are released in the BS. As such, adjustment of leaf nitrogen metabolism in C_3_-C_4_ intermediates was proposed to occur by additional metabolite shuttling between M and BS cells (Mallmann *et al*., 2014). Energy metabolism of the BS also needs to be readjusted to the additional amount of NH_3_ fixation. Depending on light availability in the M and BS cells, plants operating the glycine shuttle would need to adjust their photosynthetic pigment and protein distribution (Johnson *et al*., 2021). Shuttling of malate, aspartate, pyruvate, α-alanine, α-ketoglutarate and glutamate could contribute to rebalancing of nitrogen and energy balances between the two cell types (Mallmann *et al*., 2014; Johnson *et al*., 2021).

In the M cells, a rise in phosphoenolpyruvate carboxylase (PEPC) activity can contribute to the provision of these shuttle metabolites. PEPC fixes carbon by catalysing the addition of bicarbonate to phospho*enol*pyruvate forming the C_4_ acid oxaloacetate that is usually quickly converted into malate or aspartate. In contrast to rubisco, PEPC possesses higher substrate specificity and affinity. Combined with the decarboxylation reactions in the BS, high PEPC activity in the M cells can implement a carbon shuttle mechanism transporting C_4_ metabolites into the BS where CO_2_ is released. Plants using the glycine-in combination with such C_4_ shuttle have been identified mainly in the Asteraceae genus *Flaveria*. They are also termed as C_2_+C_4_ or C_3_-C_4_ type II species (Edwards and Ku, 1987; Sage *et al*., 2014; Bellasio, 2017).

Additional anatomic rearrangements and consequent separation of the PEPC and rubisco reactions into the M and BS cells finally support an efficient C_4_ cycle (Taniguchi *et al*., 2021). In the M cells, CO_2_ is converted into bicarbonate by carbonic anhydrase and is then fixed by PEPC. The bound carbon diffuses into the BS mainly in the form of malate or aspartate. Decarboxylation is then mediated by the NADP-malic enzyme in plastids, NAD-malic enzyme in the mitochondria or phospho*enol*pyruvate carboxykinase in the cytosol. The cycle is completed by diffusion of a C_3_ metabolite back to the M cells where ATP is needed for PEP regeneration by pyruvate phosphate dikinase. Plants with a strong C_4_ shuttle, but which still exhibit rubisco in the M are classified as C_4_-like (Moore *et al*., 1989). In *bona fide* C_4_ species, all CO_2_ is first assimilated by PEPC, with subsequent shuttling of the resulting C_4_ acid to the BS where CO_2_ is then delivered to rubisco by a decarboxylase reaction. High CO_2_ partial pressure in the BS strongly represses the oxygenase reaction, the following photorespiratory pathway and the concomitant loss of CO_2_ and NH_3_. The division of photosynthetic reaction between the two cell types however requires alterations of adjacent pathways, especially ATP and NADPH budgets, CBB cycle, nitrogen metabolism and carbon storage (Brautigam *et al*., 2011; Gowik *et al*., 2011; Schlüter and Weber, 2020).

Efficiency of the C_4_ shuttle also depends on anatomical features, especially the close connection between M and BS cells. In the majority of C_4_ species, the BS forms a tight cell layer around the veins without direct exposure to the intercellular space (Sage *et al*., 2014). The proportion of M tissue is reduced to a second cell layer around the BS cells. The interveinal distance in such Kranz anatomy is limited and C_4_ species usually have high vein densities. To prevent CO_2_ leakage, BS cell walls are often thickened and high plasmodesmata frequency at the interface between M and BS cells facilitate metabolite exchange (Danila *et al*., 2018).

Since CO_2_ fixation in C_4_ species can continue at lower internal CO_2_ concentration (Ci) and stomatal conductance, water use efficiency is improved compared to C_3_ species. Operation of rubisco under high CO_2_ partial pressure allows high efficiency for the carboxylase reaction with lower amounts of protein thus also improving the nitrogen use efficiency of C_4_ photosynthesis.

C_4_ photosynthesis evolved independently in more than 60 dicot and monocot lineages (Sage *et al*., 2011*a*) and it has been proposed that the path from C_3_ to C_4_ occurred through the above-described stages. Such an evolutionary process is supported by data from groups of closely related plants species which exhibit different CO_2_ concentrating shuttles and BS anatomy (Sage *et al*., 2012). Further support comes from modelling approaches which predict increased fitness by continuous optimisation steps between C_3_ and C_4_ (Heckmann *et al*., 2013; Mallmann *et al*., 2014; Blätke and Bräutigam, 2019; Dunning *et al*., 2019*a*). The order of adjustments could have varied in the different lineages (Williams *et al*., 2013), but installation of the initial glycine shuttle seems to be crucial (Heckmann *et al*., 2013; Blätke and Bräutigam, 2019). Depending on the preconditioning situation in the C_3_ species, the evolution to C_4_ occurred at different rates between plant lineages (Bräutigam and Gowik, 2016). Such preconditions could be related to plants genetics or environmental pressures (Christin and Osborne, 2014; Schlüter *et al*., 2017). For many C_4_ species, no close intermediate relatives have been found thus far and this could indicate that evolution from C_3_ to C_4_ happened relatively fast and smoothly in these lineages (Edwards, 2014). In *Alloteropsis semialata*, a grass that includes C_3_, C_4_ and intermediate stages, only very few transcriptional changes could be connected to the installation of a weak C_4_ cycle (Dunning *et al*., 2019*a*).

Most prominent among the glycine shuttle operating plants are the C_3_-C_4_ Brassicaceae species, a large plant family that does not contain any C_4_, but several C_3_-C_4_ intermediate species (Apel *et al*., 1997). The Brassicaceae include the model species *A. thaliana*, but also important crop species such as canola or rapeseed (*Brassica napus*), cabbage (*Brassica oleracea*), radish (*Raphanus sativus*), mustard (*Sinapis alba*) and the salad vegetable rocket or arugula (*Eruca sativa, Diplotaxis tenuifolia*). With the exception of *D. tenuifolia*, these are all C_3_ species.

The carbon concentration system of the C_3_-C_4_ intermediate species could improve leaf carbon economy during extreme weather periods that occur more frequently in the current climate conditions (Bellasio and Farquhar, 2019; Lundgren, 2020). Attempts to engineer the highly efficient C_4_ carbon concentration metabolism into rice have shown that introduction of numerous C_4_ pathway genes is possible, but introduction of the according anatomy and efficient integration of the C_4_ biochemistry into the C_3_ background is more challenging (Ermakova *et al*., 2020). Introduction of the less complex glycine shuttle could therefore be easier to achieve (Lundgren, 2020). The Brassicaceae family contains multiple species operating the photorespiratory glycine shuttle in close relationship to important crops (Schlüter *et al*., 2017). Detailed knowledge about the carbon concentration mechanisms existing in this plant group can help to identify the necessary traits for such an engineering or breeding approach.

In our study, we use the CCP to rank the species and accessions according to their carbon concentration capacity. The survey includes 28 Brassicaceae species represented by 34 accessions and focusses on the Brassiceae tribe that evolved ∼23 mya with centres of origin and diversity in the circum-Mediterranean region (Arias and Pires, 2012). Beside important crops species, this tribe includes all known C_3_-C_4_ Brassicaceae (Apel *et al*., 1997; Schlüter *et al*., 2017). So far, C_3_-C_4_ related physiological, biochemical and anatomical traits have been determined within a single origin or by comparison of few individual species. Here, our survey of multiple C_3_ and C_3_-C_4_ lineages allows us to better understand the variation in a total of 75 photosynthetic-related parameters within and between C_3_ and C_3_-C_4_ species to elucidate the characteristics which are distinctly associated with the C_3_-C_4_ photosynthesis. With this approach, we aim to identify the traits essential for the operation of carbon concentrating mechanisms within the Brassicaceae and this could guide future engineering approaches. Furthermore, we will also learn about lineage specific developments of the C_3_-C_4_ pathways and potential variation within the trait.

## Material and Methods

### Plant cultivation

The seeds were obtained from Botanical gardens, seed stock centres, and seed companies. The complete list of plant accessions included: *Arabidopsis thaliana* (At), *Brassica gravinae* (4 accessions, Bg1 to Bg4), *Brassica juncea* (Bj), *Brassica napus* (Bn), *Brassica nigra* (Bni), *Brassica oleraceae* (Bo), *Brassica rapa* (Br), *Brassica repanda* (Be), *Brassica tournefortii* (2 accessions, Bt1 and Bt2), *Carrichtera annua* (Ca), *Diplotaxis acris* (Da), *Diplotaxis erucoides* (2 accessions, De1 and De2), *Diplotaxis harra* (Dh), *Diplotaxis muralis* (Dm), *Diplotaxis tenuifolia* (Dt), *Diplotaxis tenuisiliqua* (Ds), *Diplotaxis viminea* (Dv), *Eruca sativa* (Es), *Hirschfeldia incana* (2 accessions HIR1 and HIR3), *Moricandia arvensis* (Ma), *Moricandia moricandioide*s (Mm), *Moricandia nitens* (Mn), *Moricandia sinaica* (Msi), *Moricandia spinosa* (Mp), *Moricandia suffruticosa* (Ms), *Raphanus raphanistrum* (Rr), *Raphanus sativus (*Rs) and *Sinapis alba* (Sa). From the Cleomaceae, the C_4_ species *Gynandropsis gynandra* (Gg) was also included in the present study. A complete list of origins for these seed can be found in Supplemental table S1.

Seeds were vapour sterilised by incubation in an exicator with a fresh mixture of 100ml 13% Na-Hypochloride with 3 ml of 37% HCl for 2h. The sterilised seeds were then germinated on plates containing 0.22% (w/v) Murashige Skoog medium, 50mM MES pH 5.7 and 0.8% (w/v) Agar. After 7 to 10 days, the seedlings were transferred to pots containing a mixture of sand and soil (Floraton 1 soil mixture, Floraguard, Germany) at a ratio of 1:2. All plants were firstly cultivated in climate chambers (CLF Mobilux Growbanks, Germany) under 12h day conditions with 23°C/20°C day/night temperatures and ∼200 μmol s^-1^ m^-2^ light. After establishment in soil (about two weeks), the plants were transferred to the greenhouse of the Heinrich Heine University with a 16 h day/8 h night cycle. Natural light conditions in the greenhouse were supplemented with metal halide lamps (400W, DH Licht, Germany) so that the plants received between 250 to 400 μmol m^-2^ s^-1^. Minimum temperatures were adjusted to 21°C during the night and 24°C during the day.

The initial main experiment was conducted between October 2018 and March 2019. Additional accessions were studied after the same protocol and in the same greenhouse between July and October 2020. As controls *G. gynandra, D. tenuifolia*, and *H. incana* (HIR3) were included in both experiments. Gas exchange parameters obtained for these three species, especially CO_2_ compensation points (CCP) were stable across the experiments. Thus, results from both experiments were considered comparable. Gas exchange was measured on the youngest fully expanded rosette leaves before onset of flowering. After gas exchange measurements were performed, plants were taken back to the greenhouse for two days. Following that, leaf material was harvested for metabolite analysis (only experiment in 2018/2019), EA-IRMS analysis, leaf vein determination and embedding for light microscopy. A third experiment was conducted on plant accessions selected from the initial experiments in September to November 2021 in the new greenhouses of the Heinrich-Heine University equipped with natural light LED lamps using the same experimental design. Samples for protein and PEPC assay at midday by snap freezing in liquid nitrogen. An additional leaf was used for determination of specific leaf area (SLA).

### Phylogenetic tree

Plants for genome sequencing were grown from the same seed stock in a climate chamber. Linked read sequencing was performed by 10x Genomics and BGI, complemented by PacBio and Nanopore long read sequencing for some species (Guerreiro *et al*., 2023). Most assemblies are linked read assemblies, some being scaffolded and gapfilled with the long read data, while two assemblies are long read assemblies polished and scaffolded by linked reads.

The assemblies are pseudohaploid, with alternative haplotype contigs having been removed with Purge Haplotigs (v1.1.0) (Roach *et al*., 2018). The novel genome assemblies were complemented with literature assemblies (Guerreiro *et al*., 2023). Repeat regions were identified for each assembly with Mite-hunter (Han and Wessler, 2010), genometools (v1.5.9), LTRharvest (Ellinghaus *et al*., 2008) and RepeatModeller (v1.0.11) (Smit *et al*., 2015). Those repeat regions were masked out of the assembly prior to gene annotation using RepeatMasker (v4.0.9) (Smit et al., 2015). Gene annotation was performed using Maker2 (Campbell *et al*., 2014) and the same protein database for every genome (Guerreiro *et al*., 2023). The predicted proteomes of every species were filtered for Annotation Edit Distance (AED, (Eilbeck *et al*., 2009)) values smaller than 0.5. Functional annotation was performed with AHRD in order to remove Transposon related genes (Guerreiro *et al*., 2023).

Finally, the filtered proteomes were fed into Orthofinder v2.5.1 (Emms and Kelly, 2015, 2019) for orthology identification based on all vs all sequence BLASTp searches and MCL clustering (Emms and Kelly, 2015). Multiple sequence alignments for identified orthogroups (HOGs) were produced with MAFFT and used for creating gene trees with RaxML with PROTGAMMALG substitution model. The gene trees of HOGs with single-copy genes for at least 80% of species (102 HOGs) were fed to ASTRAL-pro (Zhang *et al*., 2020) for the creation of a species phylogeny with bootstrap values for each node.

### Photosynthetic gas exchange

Gas exchange was measured on the youngest fully expanded rosette leaf about 6 to 10 weeks after sowing and before the onset of flowering. The settings of the LI6800 (LI-COR Corporate, USA) were as follows: flow of 300 μmol s^-1^, fan speed of 10,0000 rpm, light intensity of 1500 μmol m^-2^ s^-1^, leaf temperature of 25°C and VPD of 1.5 kPa. After adjustment of leaves to the conditions in the leaf chamber, A-Ci curves were measured at reference atmospheric CO_2_ concentrations of 400, 200, 100, 75, 50, 40, 30, 20, 10, 0, 400, 400, 600, 800, 1200, and 1600 ppm. In parallel, fluorescence parameters were determined for all CO_2_ conditions. For the experiment 2018/2019 the LI-6800 was equipped with a fluorescence head measuring Fv’/Fm’ and ETR at each CO_2_ level.

For calculation of the CCP and the carboxylation efficiencies (CE = initial slope of A-Ci), a minimum of four data points in the linear range close to the interception with the Ci axis were used. Maximal assimilation was determined at CO_2_ concentrations between 1200 to 1600 ppm. Assimilation (A), stomatal conductance (gsw), internal CO_2_ concentration (Ci), water use efficiency (WUE = A/gsw), and the ratio between internal and external CO_2_ concentrations (CiCa) from the measurements at 400 ppm (ambient CO_2_), 200 ppm, 100 ppm and 50 ppm CO_2_ were used for more detailed physiological analysis of the investigated plant accessions.

### Metabolite and element analysis

After the gas exchange measurements, plants were allowed to readjust to greenhouse condition before sampling for metabolite patterns. Leaves were snap frozen into liquid nitrogen directly in the greenhouse in the late morning and stored at -80°C. The leaf samples then were homogenized into a fine powder by grinding in liquid nitrogen. Soluble metabolites were extracted in a 1.5 ml extraction solution consisting of water : methanol : chloroform in a 1:2.5:1 mixture following the method of Fiehn et al. (Fiehn *et al*., 2000). 30 μl of the supernatant was dried completely in a vacuum concentrator and derivatized for GC-MS measurements (Gu, 2012). GC-MS measurements were performed as described by Shim et al. (Shim *et al*., 2020) using a 5977B GC-MSD (both Agilent Technologies). Metabolites were identified via MassHunter Qualitative (v b08.00, Agilent Technologies) by comparison of spectra to the NIST14 Mass Spectral Library (https://www.nist.gov/srd/nist-standard-reference-database-1a-v14). A standard mixture containing all target compounds at a concentration of 5 μM was processed in parallel to the samples as a response check and retention time reference. Peaks were integrated using MassHunter Quantitative (v b08.00, Agilent Technologies). For relative quantification, all metabolite peak areas were normalized to the corresponding fresh weight used for extraction and to the peak area of the internal standard ribitol or dimethylphenylalanine (Sigma-Aldrich). The same homogenised leaf material was used for determination of δ^13^C and CN ratios. After lyophilisation the material was analysed using an Isoprime 100 isotope ratio mass spectrometer coupled to an ISOTOPE cube elemental analyser (both from Elementar, Hanau, Germany) according to (Gowik *et al*., 2011). In most cases, a minimum of four biological replicates per plant accessions were analysed.

### Vein density measurements

Mature rosette leaves were used for vein density measurements. The leaf material was cleared in an acetic acid: ethanol mix (1:3) overnight followed by staining of cell walls in 5% safranin O in ethanol, and de-staining in 70% ethanol. Pictures were taken using a Nikon eclipse Ti-U microscope equipped with a ProgRes MF camera from Jenoptik, Germany, at 4x magnification. The vein density was analysed with ImageJ software. In most cases, six leaves were analysed for vein density per line with a minimum of three pictures measured and averaged per leaf.

### Specific leaf area (SLA)

Whole mature rosette leaves were cut and their outlines were copied on checked paper. The fresh weight (FW) was taken immediately after that. The dry weight (DW) was determined after 48 h at 60°C. The leaf area was determined using ImageJ software. For calculation of SLA, the area was divided by the dry weight.

### Analysis of leaf cross section

For light microscopy, sections of ca. 1 × 2 mm were cut from the top third of mature rosette leaves and immediately fixed in 2% paraformaldehyde, 2% glutaraldehyde, 0.1% Triton-X-100 in phosphate buffer saline (137 mM NaCl, 2.7mM KCl, 12mM H_2_PO_4-_/HPO_42-_, pH 7.4). Vacuum was applied to the reaction tubes until all leaf sections sank to the bottom. The sections were incubated in the primary fixation solution overnight followed by washing once with phosphate buffer saline solution, pH 7.4, and twice with distilled H_2_O. For post-fixation, the sections were incubated in 1% OsO_4_ for 45 min followed again by washing three times with distilled H_2_O. A dehydration series was performed ranging from 30% to 100% acetone, followed by incubation in increasing proportions of Araldite resin until 100% Araldite was reached. The sections were finally positioned into flat embedding moulds and polymerised at 65°C for at 48h.

After cutting, the leaf sections were stained with toluidine blue solution (0.5% toluidine blue, 0.5% methylene blue, 6% Na_2_B_4_O_7_, 1% H_3_BO_3_) and studied under a light microscope (Zeiss Axio Observer, Carl Zeiss, Germany). For quantitative analysis, pictures of at least three BS per biological replicate were taken and analysed with ImageJ software. The following parameter were determined per BS for quantitative analysis: cross section area of BS cell (BS_cell_area), area of organelles orientated towards the vein (V_organelle_area), area of organelles orientated towards intercellular space (ICS) and M (M_organelle_area). The following parameter were calculated: percent of vein orientated organelle area per BS cell area (percent_V_organelle), percent of organelles orientated towards ICS and M (percent_M_organelles), the total organelle area per cell (V_organelle_area + M_organelle_area = total_organelle_area) and the ratio between percent values for vein and ICS/M orientated organelles. Furthermore, the leaf thickness was measured at the site of the selected BS. Three representative cells were analysed per BS, and three BS were analysed per plant.

### PEPC activity

Total soluble proteins were extracted from the homogenised leaf material in 50mM Hepes-KOH pH 7.5, 5mM MgCl2, 2mM DTT, 1mM EDTA, 0.5% Triton-X-100. For the PEPC assay 10 μl of the extract were mixed with assay buffer consisting of 100mM Tricine-KOH pH 8.0, 5mM MgCl_2_, 2mM DTT, 1mM KHCO_3_, 0.2mM NADH, 5mM glucose-6-phosphate, 2U ml^-1^ malate dehydrogenase in a microtiter plate. The reaction was started after addition of phosphoenolpyruvate to a final concentration of 5mM in the assay. Protein content of the solutions was determined with the BCA assay (Thermo Fischer Scientific).

### Statistical analysis

Data analysis was performed using R (www.R-project.org). Statistical differences between the measured parameters in the accessions were calculated by one-way ANOVA followed by Tukey post-hoc test. Differences between parameters in C_3_ and C_3_-C_4_ photosynthesis types were determined by a two-tailed t-test (Supplemental figures S4, S5, S6). Geographical distribution data was downloaded from the gbif.org website (August 2022), associated climate data was retrieved from the WorlCLim version 2.1 database (WorldClim.org) using 10 min resolution.

## Results

### Assessment of CO_2_ concentrating efficiency by measuring CO_2_ compensation points

The CCP is a measure of the internal leaf CO_2_ level at which photosynthetic CO_2_ fixation is equal to the CO_2_ release by photorespiration, day respiration and other catalytic processes (i.e., the concentration at which net CO_2_ assimilation is zero). In the present study, the CCP was determined from A-Ci curves across a diverse range of 34 Brassicaceae accessions from 28 species to assess their carbon usage efficiency (Figure 1). The closest known C_4_ relative in the Cleomaceae, *G. gynandra*, was also included in the analysis for comparison.

**Figure 1.**
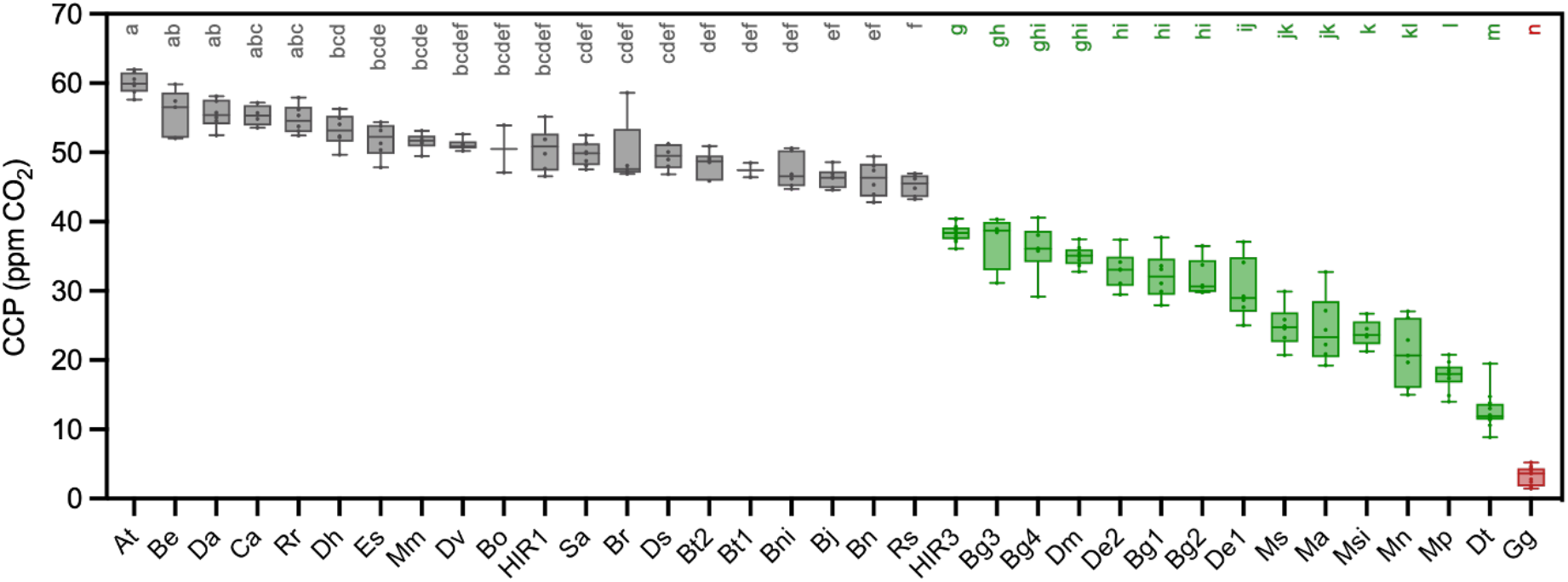
CO_2_ compensation points in selected Brassicaceae species. CO_2_ compensation points were measured in young, fully expended leaves of greenhouse grown plants. The letters above each box indicate the statistical grouping determined by ANOVA followed by HSD post-hoc test with a=0.05. The tested plant lines are colored according to photosynthesis type as C_3_ (grey), C_3_-C_4_ (green) and C_4_ (red). Species and accessions have been abbreviated for legibility and are provided in full in Material and Methods.

When sorting all sampled plant accessions in the Brassicaceae according to their CCP, a range of CCP values from 60 to 10 ppm was detected. The Cleomaceae C_4_ species *G. gynandra* with highly efficient CO_2_ concentration showed CCP values < 10 ppm. Plants with CCP values between 10 and 40 ppm are predicted to utilize less efficient CO_2_ concentration, such as the photorespiratory shuttle, and were hereby classified as C_3_-C_4_ intermediates. In contrast, all accessions and species with a CCP >40 ppm were classified as C_3_ species. This grouping was supported by ANOVA with post-hoc Tukey HSD test (alpha = 0.05; Figure 1). The same threshold values were also used in the survey by Krenzer et al. (Krenzer *et al*., 1975).

In our study, *A. thaliana* exhibited the highest CCP of 60.1 ppm. Slightly lower CCPs between 45 and 60 ppm were observed for the C_3_ species of the Brassiceae clade, including *Brassica repanda, Diplotaxis acris, Carrichtera annua, Raphanus raphanistrum, Diplotaxis harra, Eruca sativa, Moricandia moricandioides, Diplotaxis viminea, Brassica oleraceae, Hirschfeldia incana* accession HIR1, *Sinapis alba, Brassica rapa, Diplotaxis tenuisiliqua, Brassica tournefortii, Brassica nigra, Brassica juncea, Brassica napus* and *Raphanus sativus*. In comparison to these C_3_ plants, a significant reduction in the CCPs was observed in 14 accessions classified in the present study as C_3_-C_4_ intermediates, these included *Hirschfeldia incana* accession HIR3, four accessions of *Brassica gravinae, Diplotaxis muralis*, two accessions of *Diplotaxis erucoides, Moricandia suffruticosa, Moricandia arvensis, Moricandia sinaica, Moricandia nitens, Moricandia spinosa* and *Diplotaxis tenuifolia*.

Among the C_3_-C_4_ intermediates, the lowest CCP value of 12 ppm was measured in *D. tenuifolia*. In contrast, the highest CCP value recorded among the C_3_-C_4_ intermediates at ∼40 ppm was measured in *H. incana* (accession HIR3). Importantly, the identification of *H. incana* HIR3 as a C_3_-C_4_ intermediate species which operates a CO_2_ concentrating mechanism is described here for the first time. Interestingly, another closely related *H. incana* accession (HIR1) exhibited a CCP value within the range of C_3_ species (Figure1). In all other species different accessions were assigned to the same photosynthetic type. Altogether, the wide range of CCPs among Brassicaceae and especially the C_3_-C_4_ intermediates indicates that the underlying biochemical and anatomical mechanisms are not necessarily conserved in this clade and could be connected to independent evolutionary origins.

### Phylogeny suggests up to five independent origins of C_3_-C_4_ photosynthesis in the Brassiceae

The phylogenetic relationship among the plant species selected for this study was investigated using sequence data from 102 orthogroups (Figure 2). When investigating the distribution of species classified as C_3_-C_4_ intermediates based on CCP data, the tree reveals multiple origins of CO_2_ concentration mechanisms in the Brassiceae. We note that the presented tree includes only a small subset of species from the Brassiceae group, though a recently published and more densely sampled phylogenetic tree (Koch and Lemmel, 2019) supports these claims.

**Figure 2.**
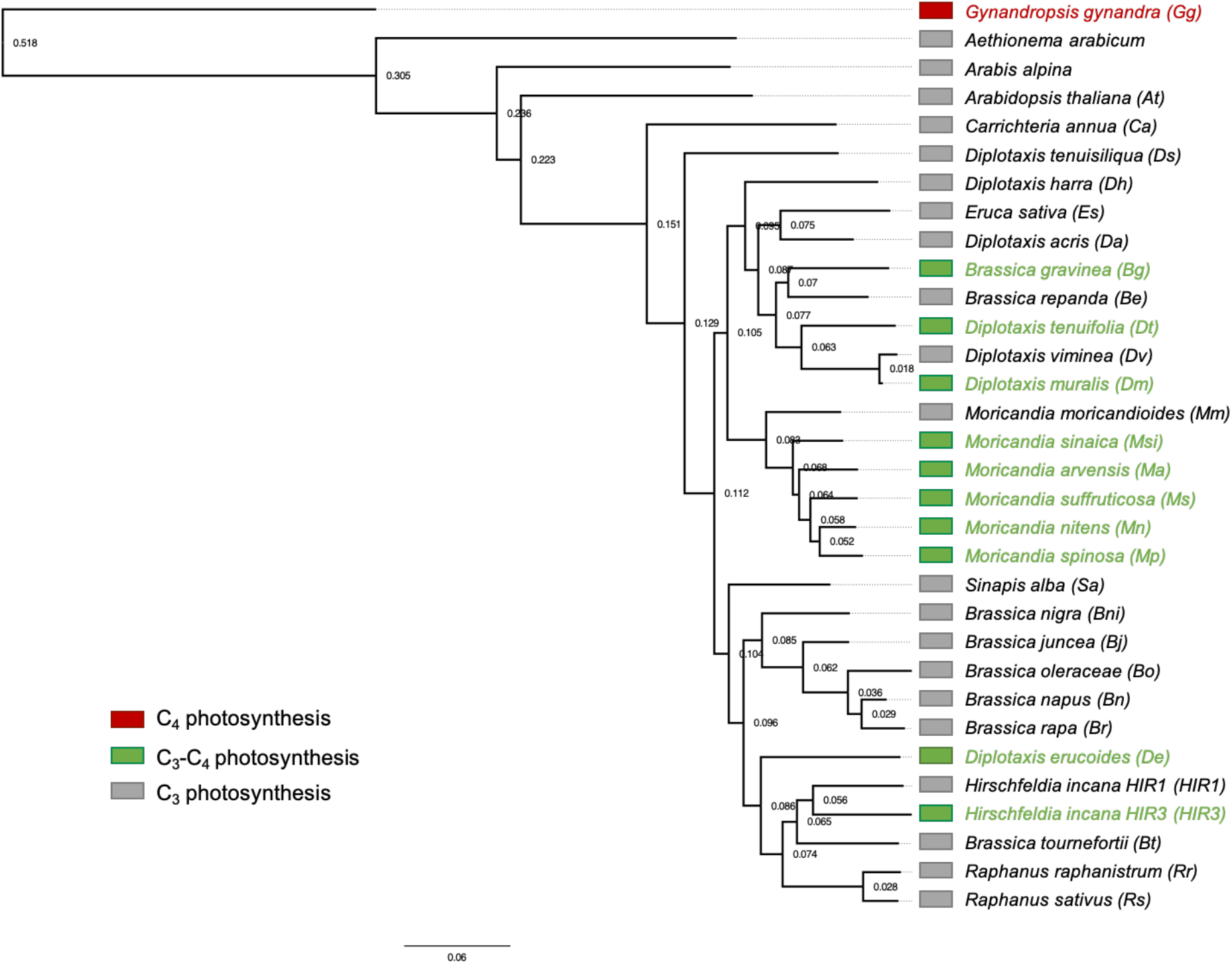
Phylogeny and photosynthesis types. The number on the nodes represents bootstrap values. Species abbreviations are given in brackets. Plant species and accessions are colored according to the photosynthesis type as C_3_ (grey), C_3_-C_4_ (green) and C_4_ (red).

The phylogenetic positions of *B. gravinae*, the *H. incana* accession HIR3 and *D. erucoides* on the tree suggest independent origins of C_3_-C_4_ features. Moreover, in the *Moricandia* group, C_3_-C_4_ features are observed among the close relatives *M. arvensis, M. suffruticosa, M. nitens, M. sinaica* and *M. spinosa*, but not the sister species *M. moricandioides* which is C_3_. Thus, C_3_-C_4_ most likely evolved once in this group, in a common ancestor preceding the speciation of *M. arvensis, M. suffruticosa, M. nitens, M. sinaica* and *M. spinosa* but after diversification from *M. moricandioides*. Finally, C_3_-C_4_ like CCPs were observed in *D. tenuifolia* and *D. muralis*. Both of these respective species are closely related to the C_3_ species *D. viminea. D. muralis* is a natural hybrid between the C_3_ parent *D. viminea* and the C_3_-C_4_ parent *D. tenuifolia*. Therefore, the C_3_-C_4_ features in *D. muralis* are assumed to be inherited from *D. tenuifolia* (Ueno *et al*., 2006). In summary, our phylogenetic data indicate that C_3_-C_4_ features have independently evolved up to five times in the Brassicaceae, in *B. gravinae*, in *D. erucoides*, in *H. incana*, in the *Moricandia* group and in *D. tenuifolia* (subsequently inherited by hybridisation in *D. muralis*).

### Physiology of C_3_, C_3_-C_4_ and C_4_ leaves under different CO_2_ concentration

Efficiency of photosynthetic gas exchange in the different C_3_ and C_3_-C_4_ Brassicaceae accessions was assessed under ambient CO_2_ (400 ppm) and saturating light (1500 μmol m^-2^ s^-1^). Here, under ambient conditions, no association between photosynthesis type and net assimilation could be observed (Figure 2 A). For instance, in C_3_ plants, assimilation rates under ambient CO_2_ varied between 12.3 μmol m^-2^ s^-1^ (*A. thaliana*) and 28.1 μmol m^-2^ s^-1^ (*D. tenuisiliqua*) (Figure 2A, Supplemental table S2), whilst among C_3_-C_4_ intermediates, assimilation rates varied between 17.3 μmol m^-2^ s^-1^ (*B. gravinae* accession 2) and 26.8 μmol m^-2^ s^-1^ (*D. erucoides* accession 1). Moreover, assimilation rates achieved in the C_4_ species *G. gynandra* of 23.7 μmol m^-2^ s^-1^ were similar to rates in non-C_4_ plants. Thus, assimilatory capacity under ambient CO_2_ conditions appears to be species- or accession-specific, rather than determined by plant photosynthetic machinery.

In contrast to the above, enhanced rates of assimilation were discovered in plants operating a CO_2_ concentrating mechanism under lower atmospheric CO_2_ concentrations (Figure 3C, E,Supplemental table S2). For instance, at pre-industrial levels of 200 ppm CO_2_, the C_4_ *G. gynandra* showed higher assimilatory capacity compared to any other C_3_ or C_3_-C_4_ species (Figure 3C; Supplemental table S2). Further, this elevated assimilation rate observed in *G. gynandra* became even more pronounced under 100 ppm CO_2_ (Figure 3E; Supplemental table S2). Interestingly, C_3_-C_4_ intermediate species tended to performed better than C_3_ plants under sub-ambient CO_2_ conditions (Figure 3C, E; Supplemental table S2). On average across all plants of each photosynthesis type, assimilation was significantly higher in the C_3_-C_4_ group compared to the C_3_ group under CO_2_ conditions of 200 ppm and below (t-test, p<0.05; Supplemental figure S4). Thus, although CO_2_ concentrating mechanisms do not improve net assimilation under present atmospheric CO_2_, they appear to be advantageous under former pre-industrial levels of CO_2_.

**Figure 3.**
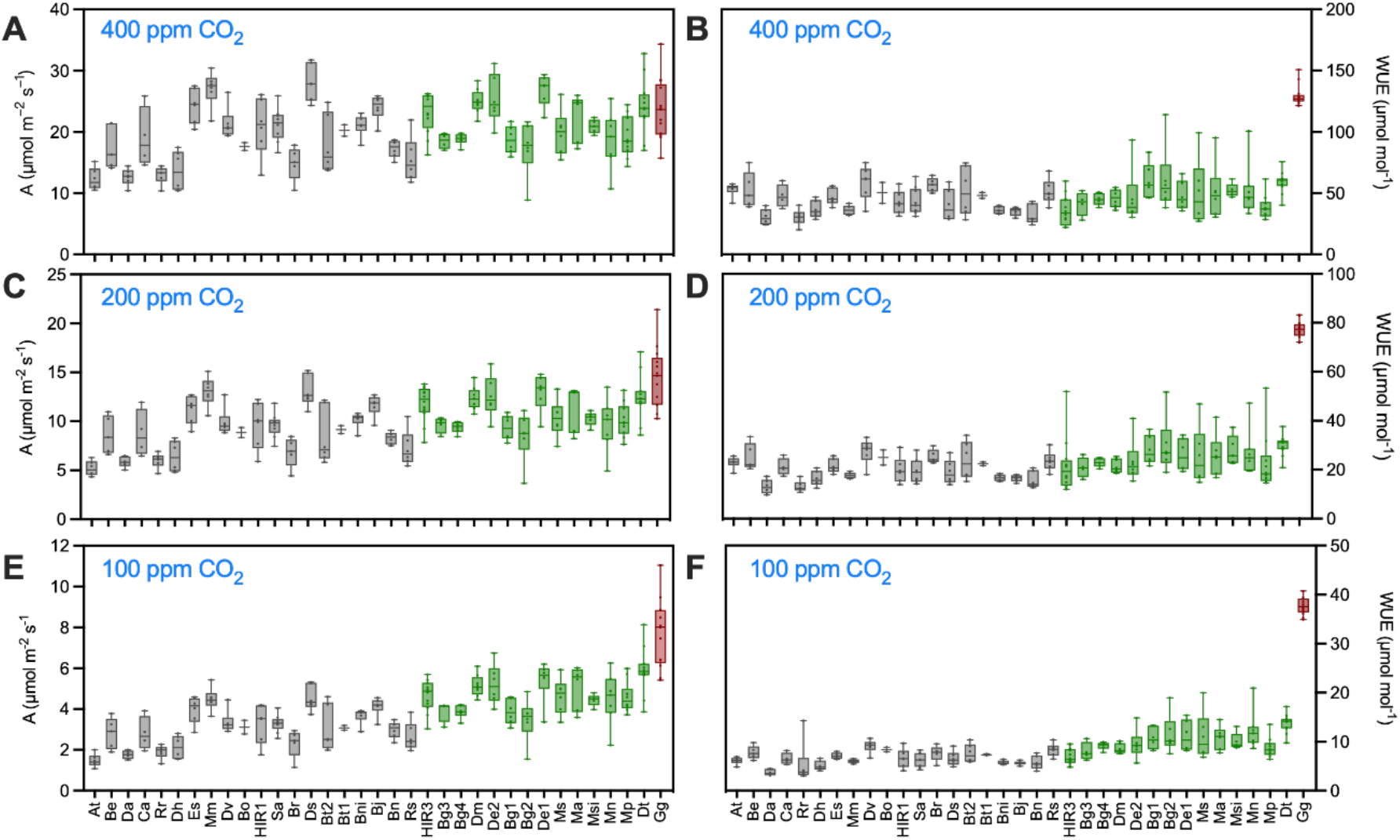
Net assimilations (A) and Water use efficiency (WUE) in under different CO_2_ concentrations. Assimilation (A, C, E) was determined in a CO_2_ response curve. WUE (B, D, F) was calculated as ratio between assimilation (A) and stomatal conductance (gsw). Gas exchange parameters were measured under conditions of ambient CO_2_ (A, B, 400 ppm) or reduced CO_2_ concentrations (C, D, 200 ppm; E, F, 100 ppm). The accessions were sorted according to their CO_2_ compensation points and colored according to the photosynthesis type as C_3_ (grey), C_3_-C_4_ (green) and C_4_ (red). Species and accessions have been abbreviated for legibility and are provided in Figure 2 and Material and Methods.

In addition to the above, operating a CO_2_ concentrating mechanism also yielded benefits in terms of improved water-use efficiency (WUE; ratio between CO_2_ assimilation and water stomatal conductance). For example, under ambient 400 ppm CO_2_, the WUE was significantly higher in the C_4_ species *G. gynandra* as compared to all other C_3_ and C_3_-C_4_ species (Figure 3 B). On average, WUE did not differ between C_3_ and C_3_-C_4_ accessions at 400 ppm CO_2_. However, C_3_-C_4_ plants were found to exhibit a significantly improved WUE at both 200 and 100 ppm CO_2_ compared to C_3_ species (t-test, p>0.05; Supplemental figure S4) recapitulating the trend observed for assimilation rate. In addition, a strong negative correlation between WUE and Ci was found across species at all atmospheric CO_2_ concentrations (Supplemental figure S1). Given that changes in stomatal conductance exhibited no photosynthesis type specific pattern (Supplemental figure S4), this result suggests that higher WUE is achieved across species in the Brassicaceae by CO_2_ assimilation at lower internal CO_2_ concentration (Ci), Thus, C_3_-C_4_ species are able to operate at lower internal CO_2_ concentrations than species from the C_3_ group.

Next, to determine the effect of C_3_, C_3_-C_4_ and C_4_ metabolism on the light reactions of photosynthesis, chlorophyll fluorescence parameters were also measured. In general, a positive correlation was found between net assimilation and electron transport rate (ETR), as well between assimilation rate and effective quantum efficiency Fv’/Fm’ (Supplemental Figure S1). However, fluorescence parameters were less affected under reduced atmospheric CO_2_ concentrations as compared to assimilation rate (Supplemental figure S1). For instance, no correlation effect was observed between CCP and ETR or Fv’/Fm’ under any CO_2_ concentration (Figure 4 C, D). Though, it should be noted that it is possible that effects on the light reactions would become obvious only after longer adjustment periods to changes in CO_2_ environmental conditions which were not able to be accommodated in the current experimental setup.

**Figure 4.**
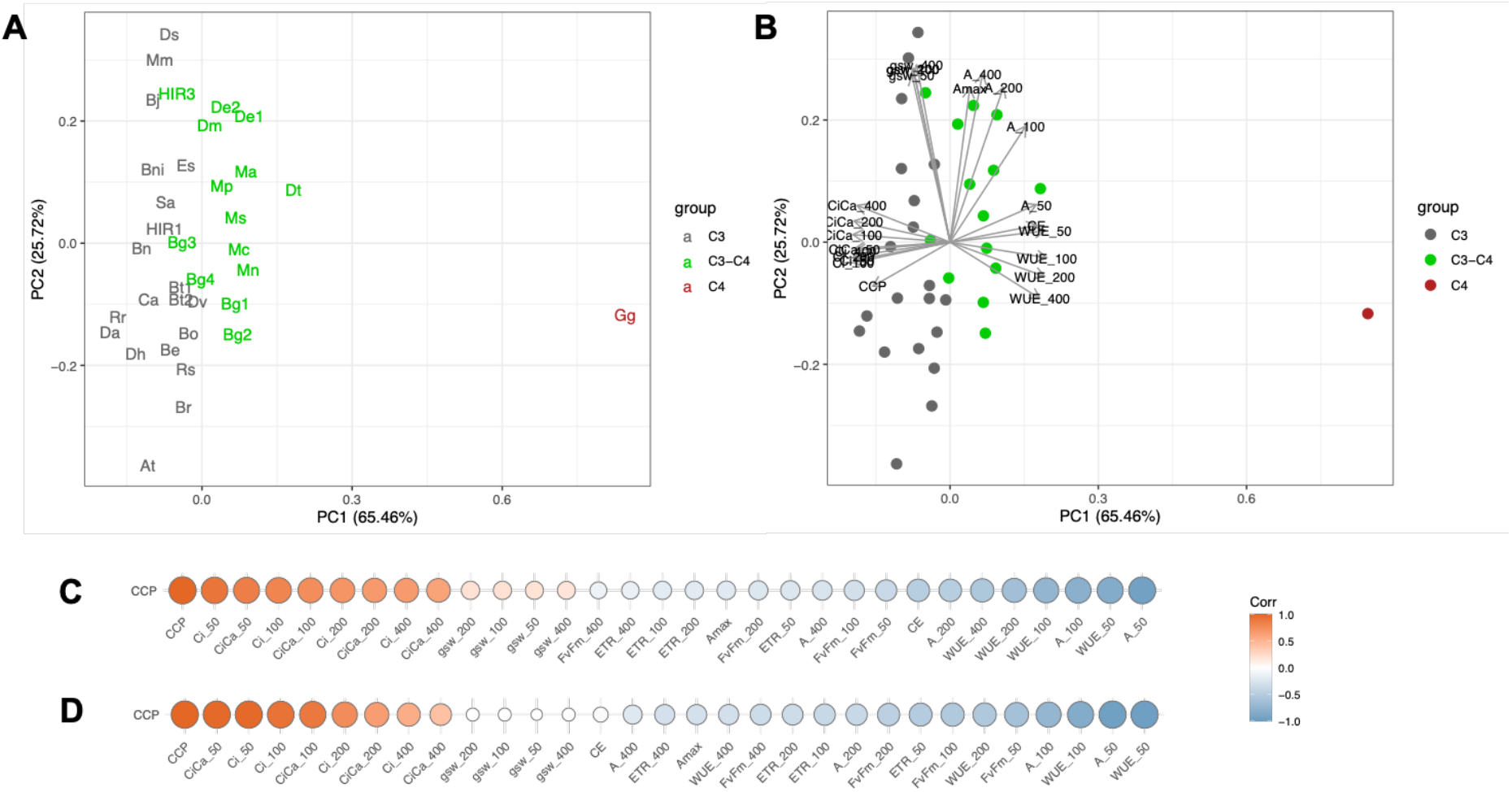
Principal component analysis (PCA) and correlations of the CO_2_ compensation point (CCP) with selected gas exchange parameters. Average values for the selected photosynthetic parameters determined under 50 to 400 ppm of CO_2_ were used for the analysis. (A) Localization of the plant lines in the PCA, (B) PCA including parameter loadings, (C) Pearson correlation coefficients demonstrated as heatmaps using all plant lines, (D) Pearson correlation coefficients demonstrated as heatmaps using only C_3_ and C_3_-C_4_ lines. The tested plant lines were colored according to photosynthesis types as C_3_ (grey), C_3_-C_4_ (green) and C_4_ (red). Species and accessions have been abbreviated for legibility and are provided in Figure 2 and Material and Methods. The numbers after the parameter abbreviation indicate the CO_2_ concentration in the outside the leaf in the measuring cuvette.

To further assess how different photosynthesis types are characterised by differences in leaf physiological parameters, a principal component analysis (PCA) was performed. Since the importance of carbon concentrating mechanisms becomes more obvious when CO_2_ is limited, gas exchange measurements at CO_2_ concentrations of 200, 100 and 50 ppm were included in this analysis in addition to those measured under ambient 400 ppm CO_2_ (Figure 4 A, B). In this PCA, the first principal component was found to explain 65.5% of the variation, and separates the C_4_ species *G. gynandra* from all other C_3_ and C_3_-C_4_ plants. To a lesser extent, the same component also separates C_3_ and C_3_-C_4_ plants though these groups do overlap on this axis (Figure 4 A, B). As expected from the above results, the first principal component is driven by WUE, Ci, CiCa ratio, CCP and carboxylation efficiency (CE). In contrast, the second principal component, driven by stomatal conductance and assimilation at higher CO_2_ concentrations, has no effect on separating plants across different photosynthesis types and is driven by species-specific variation.

As CCP was used above to classify different photosynthesis types, correlations were investigated between CCP values across species and all other measured leaf physiological parameters. A strong positive correlation was observed between CCP and both Ci and Ci/Ca ratio at low CO_2_ concentrations, respectively (Figure 4 C, D). In contrast, a negative correlation was found between CCP and both WUE and assimilation at low CO_2_ (Figure 4 C, D). These relationships were independently observed irrespective of whether the C_4_ species *G. gynandra* was included in the analysis or not. Conversely, carboxylation efficiency (CE = initial slope of A-Ci curve) was negatively correlated with CCP only when the C_4_ *G. gynandra* was included in the analysis (Figure 4C). This indicates that CE was not influenced considerably during the transition from C_3_ to C_3_-C_4_, but only during transition from C_3_-C_4_ to C_4_.

### Metabolite profiles of leaves with different photosynthesis pathways

To assess the identity of potential transport metabolites used in C_3_-C_4_ intermediates, leaf primary metabolites of sampled species were also quantified and analysed by PCA (Figure 5 A, B). In this analysis, the first principal component explained 28.25% of variation and distinguishes C_3_/C_3_-C_4_ and C_4_ leaf biochemistry. Mainly responsible for this separation are high levels of α-alanine, α-ketoglutarate, aspartate, glycine, glutamate, pyruvate, phenylalanine and γ-aminobutyric acid (GABA) in the C_4_ *G. gynandra* compared to the C_3_ and C_3_-C_4_ background (Figures 5 B; Supplemental table S3). In contrast, the second principal component sorts the majority of C_3_ species (clustered to the top of this axis) from C_3_-C_4_ species (clustered to the bottom of this axis) (Figure 5 A, B). Here, on this axis, the C_3_-C_4_ plants tend to have higher levels of serine, branched amino acids and proline, whilst the C_3_ species are characterised by higher levels of glucose, sucrose and myo-inositol.

**Figure 5.**
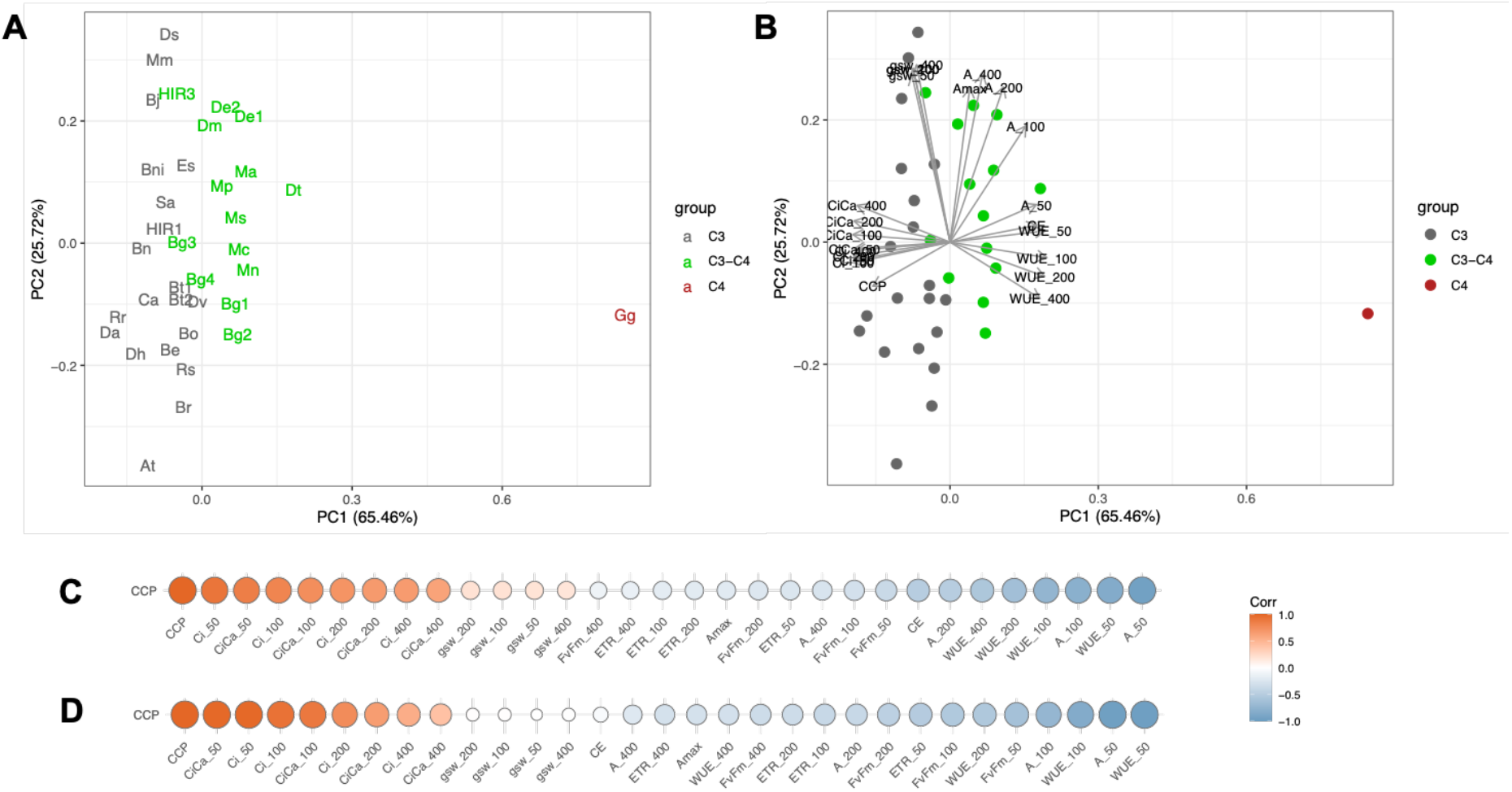
Principal component analysis (PCA) and correlations of the CO_2_ compensation point (CCP) with specific metabolites. Average values for the selected metabolites per line were used for the analysis. (A) Localization of the plant lines in the PCA, (B) PCA including metabolite loadings, (C) Pearson correlation coefficients demonstrated as heatmaps using all plant lines, (D) Pearson correlation coefficients demonstrated as heatmaps using only C_3_ and C_3_-C_4_ lines. The tested plant lines were colored according to photosynthesis type as C_3_ (grey), C_3_-C_4_ (green) and C_4_ (red). Species and accessions have been abbreviated for legibility and are provided in Figure 2 and Material and Methods.

Correlation analyses of CCP values with primary metabolite levels were also performed across species (Figure 5 C, D, Supplemental figure S2). A negative correlation was observed between CCP and the C_4_ related metabolites α-alanine, α-ketoglutarate, aspartate, glutamate, pyruvate (Figures 5, 6; Supplemental table S3). However, this relationship seems to be driven by the strong accumulation of these metabolites in the C_4_ species alone (Figure 5 C). To identify metabolites that are potentially specific to C_3_-C_4_ photosynthesis, the correlation analysis was repeated without the C_4_ outgroup species. This resulted in the reduction of the strength of all statistical associations. Specifically, only serine and glycine showed significant negative correlations with CCP (Figure 5 D; Supplemental table S3). Thus, this suggests that serine and glycine have a ubiquitous role in the glycine shuttle across all C_3_-C_4_ intermediates in the Brassicaceae. Interestingly, however, glycine was the only metabolite that increased between C_3_ and C_3_-C_4_ species which was also high in the C_4_ species (Figure 6 A; Supplemental figures S2, S5). In contrast, serine was enhanced among C_3_-C_4_ species compared to C_3_ species, but was detected at a C_3_ level in the C_4_ *G. gynandra* (Figure 6 B; Supplemental figures S2, S5). In the present study, glutamate, α-alanine, aspartate, pyruvate, malate and α-ketoglutarate formerly predicted to be involved in nitrogen shuttling of C_3_-C_4_ leaves (Mallmann *et al*., 2014) were not associated with CCP (Figure 6). Instead, levels of these metabolites were high in only some C_3_-C_4_ accessions. For instance, glutamate and aspartate levels were relatively high in *M. arvensis, D. muralis* and *D. tenuifolia*, but not in the other C_3_-C_4_ *Moricandia* species *M. nitens* and *M. suffruticosa* (Figure 6). In contrast, the two *D. erucoides* accessions separated from the majority of other C_3_-C_4_ species in the PCA (Figure 5 A, B) showed relatively high levels of glycerate, glycolate and malate (Figure 6; Supplemental table S3). In summary, the present results describe a general role of only glycine and serine as predicted shuttle metabolites in C_3_-C_4_ biochemistry across all species. However, individual biochemical adjustments are possibly active among different C_3_-C_4_ lineages.

**Figure 6.**
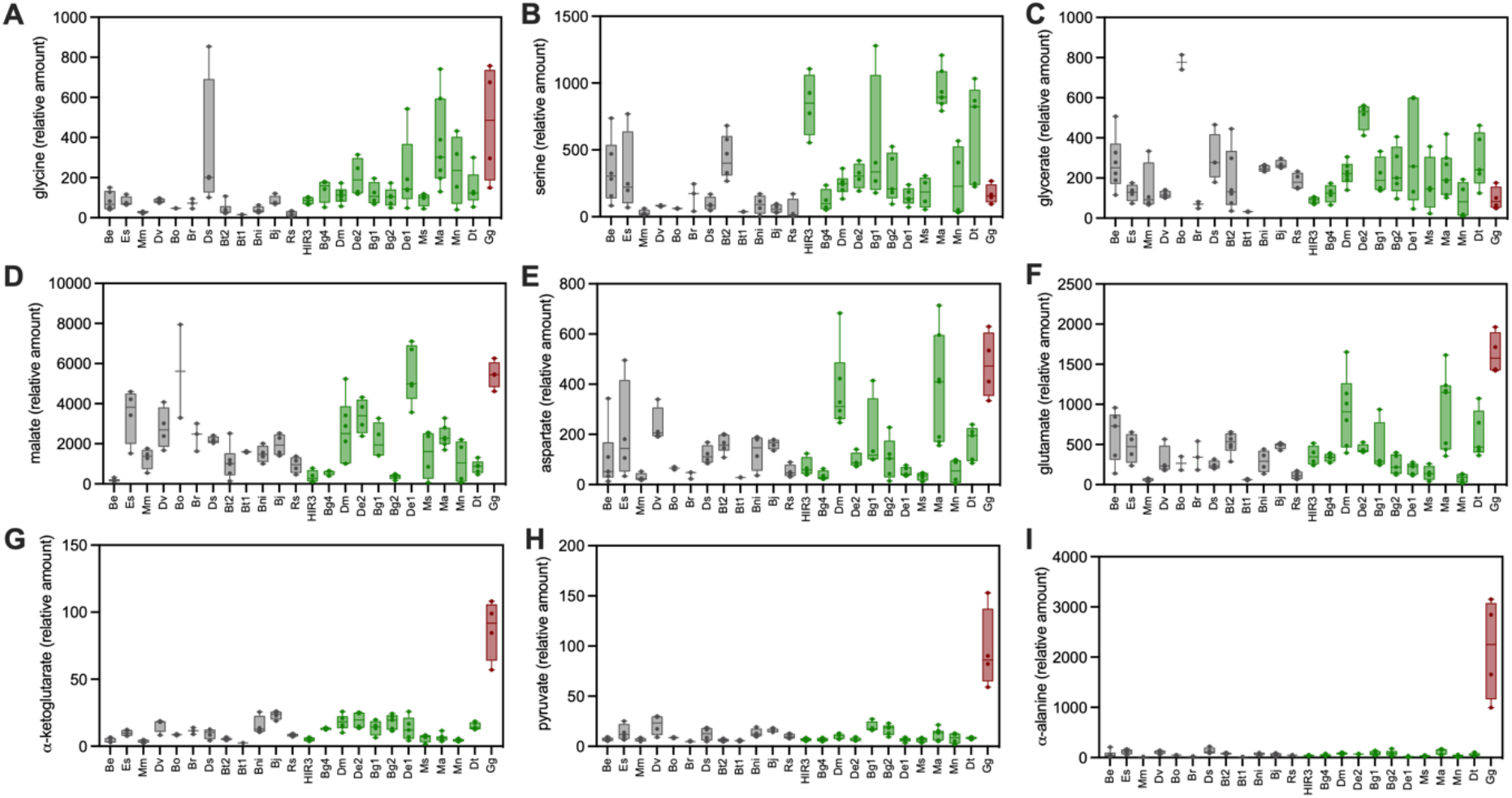
Selected Metabolites in mature leaves. Relative amounts of glycine (A), serine (B), glycerate (C), malate (D), aspartate (E), glutamate (F), D-ketoglutarate (G), pyruvate and D-alanine in selected plant accessions. The accessions were sorted according to their CO_2_ compensation point and colored by photosynthesis type (grey = C_3_, green = C_3_-C_4_ and red = C_4_). Species and accessions have been abbreviated for legibility and are provided in Figure 2 and Material and Methods.

### Association of CCP with structural features of bundle sheath

Given that leaf and BS cell architecture play an important role in underpinning CO_2_ concentrating mechanisms by enabling adequate metabolite transport between M and BS tissue, we also sought to characterise the leaf anatomy of our Brassicaceae species. In the present study, it was observed that vein density was highest in the C_4_ *G. gynandra* compared to all other species. However, no difference in vein density was observed between C_3_ and C_3_-C_4_ plant accessions (Figure 7 A). To determine whether differences in BS structure were present between photosynthetic types, a representative subset of plant accessions were studied in more detail by light microscopy (Supplemental figure S7). In this analysis, although BS cross section area was high in the C_4_ species as well as several C_3_-C_4_ species, it was not found to be significantly different between C_3_ and C_3_-C_4_ plants (Figure 7 B, Supplemental figure S6). Moreover, within the BS cells, the areas occupied by plastids and other organelles with either vein (inner half) or intercellular space (ICS)/M orientation (outer half) were determined. Areas with ICS/M oriented organelles did not differ between C_3_ and C_3_-C_4_ leaf cross section (Figure 7E). In the C_4_ leaf, none of the BS organelles faced the outer ICS/M side. On the other hand, all plant accessions with a CCP below 40 ppm featured enhanced organelle accumulation around the vein (Figure 7 H). This resulted in higher total organelle area in the C_3_-C_4_ BS cells. Thus, organelle abundance and orientation likely played a decisive role for the functioning of weak CO_2_ concentrating mechanisms.

**Figure 7.**
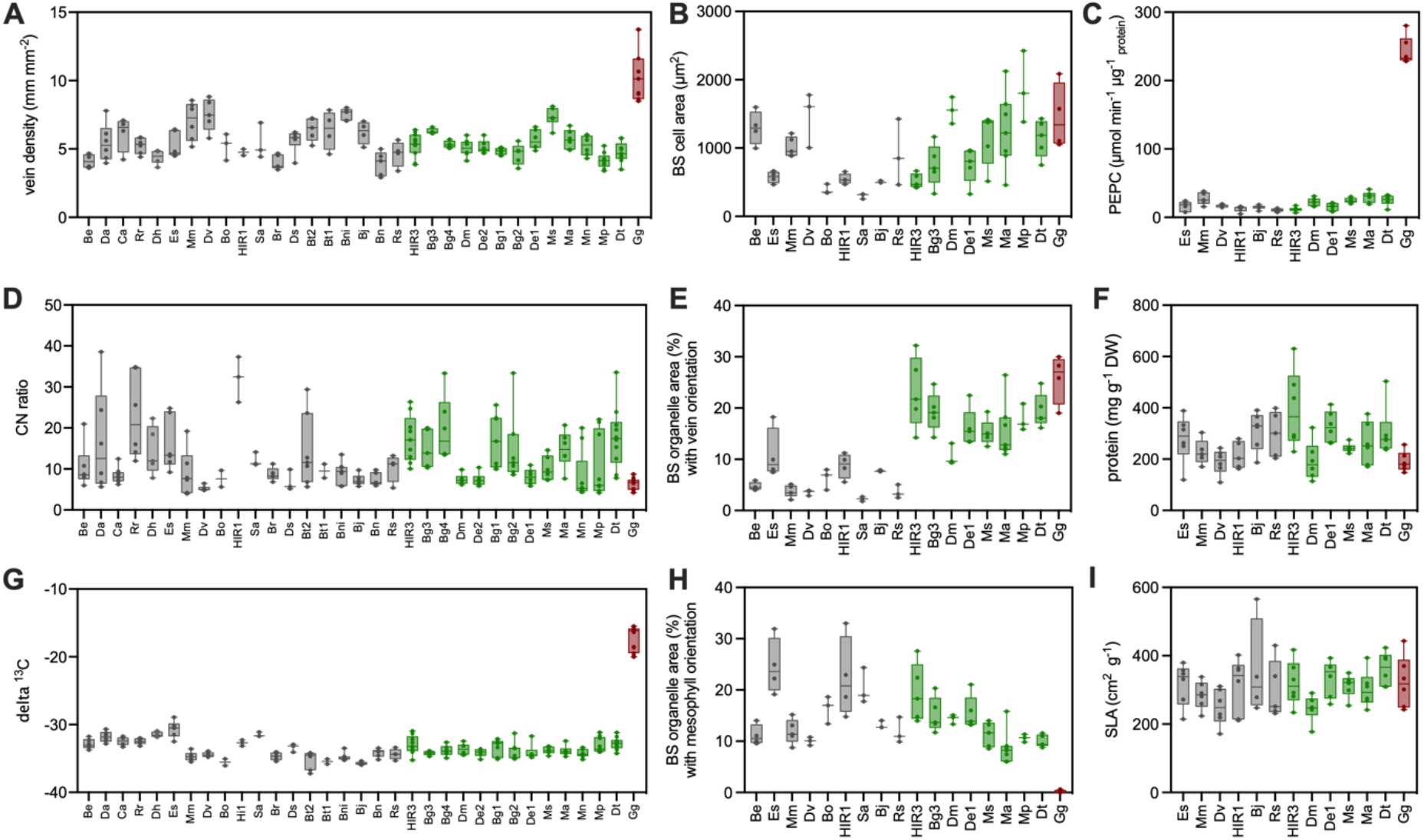
Leaf structure and composition related parameters and PEPC activity. Mature leaves were used for determination of vein density (A), bundle sheath cell area in micrographs (B), PEPC activity (C), carbon to nitrogen ratio (D), area of bundle sheath organelles with vein orientation in micrographs (E), protein content (F), ^13^C signature (G), area of bundle sheath organelles with orientation to intercellular space or mesophyll in micrographs (H), and specific leaf area (I). The accessions were sorted according to their CO_2_ compensation point and colored to photosynthesis type (grey = C_3_, green = C_3_-C_4_ and red = C_4_). Species and accessions have been abbreviated for legibility and are provided in Figure 2 and Material and Methods.

C_4_ anatomy consists of just one layer of BS and M cells around the veins which limits the total number of cell layers. In our study, the C_4_ leaves of *G. gynandra* were comparably thin. However, leaf thickness within the C_3_ and C_3_-C_4_ groups showed species-specific variation. For instance, independent of photosynthesis type, all *Moricandia* species possessed thick succulent leaves. Values of SLA (specific leaf area = area per g dry weight) are usually greater for C_4_ than for C3 leaves (Atkinson *et al*., 2016), but no pronounced photosynthesis type related differences in SLA could be observed in our study (Figure 7 I).

The C_4_ pathway allows plants to fix CO_2_ with lower nitrogen input. This means that typically, C_4_ plants have lower leaf nitrogen concentrations compared to C_3_ species (Long, 1999; Craine *et al*., 2005; Gowik *et al*., 2011). Interestingly, however, C_4_ *G. gynandra* in our analysis had surprisingly high leaf nitrogen concentrations and low leaf CN ratios (Figure 7 D). This result could be influenced by the slow growth rate of this species in comparison with the majority of Brassicaceae species in this study. However, leaf protein concentrations in this C_4_ *G. gynandra* were low relative to the background of other species indicating C_4_-specific differences in N allocation (Figure 7 F). Interestingly, no difference in CN ratios or leaf protein concentrations could be observed between the C_3_ and C_3_-C_4_ species (Figure 7 D; Supplemental figure S6).

Operation of the C_4_ pathway required increased activity of PEPC, but allows reduction in concentrations of rubisco and CBB cycle enzymes (Brautigam *et al*., 2011; Gowik *et al*., 2011). In our study, PEPC activity was 8 to 20-fold higher in the C_4_ *G. gynandra* leaves as compared to the leaves of the C_3_ and C_3_-C_4_ species (Figure 7 C, Supplemental table S4). PEPC activities varied in the individual plant accessions, but were not significantly different between the C_3_ and C_3_-C_4_ groups (Supplemental figure S6). Especially *E. sativa* and *M. moricandioides* showed PEPC activities similar to the C_3_-C_4_ accessions *M. arvensis, M. suffruticosa* and *D. tenuifolia* (Figure 8, Supplemental figure S6). These results emphasise the power of our multi-species analysis that allows distinction between species and photosynthesis type related variation.

**Figure 8.**
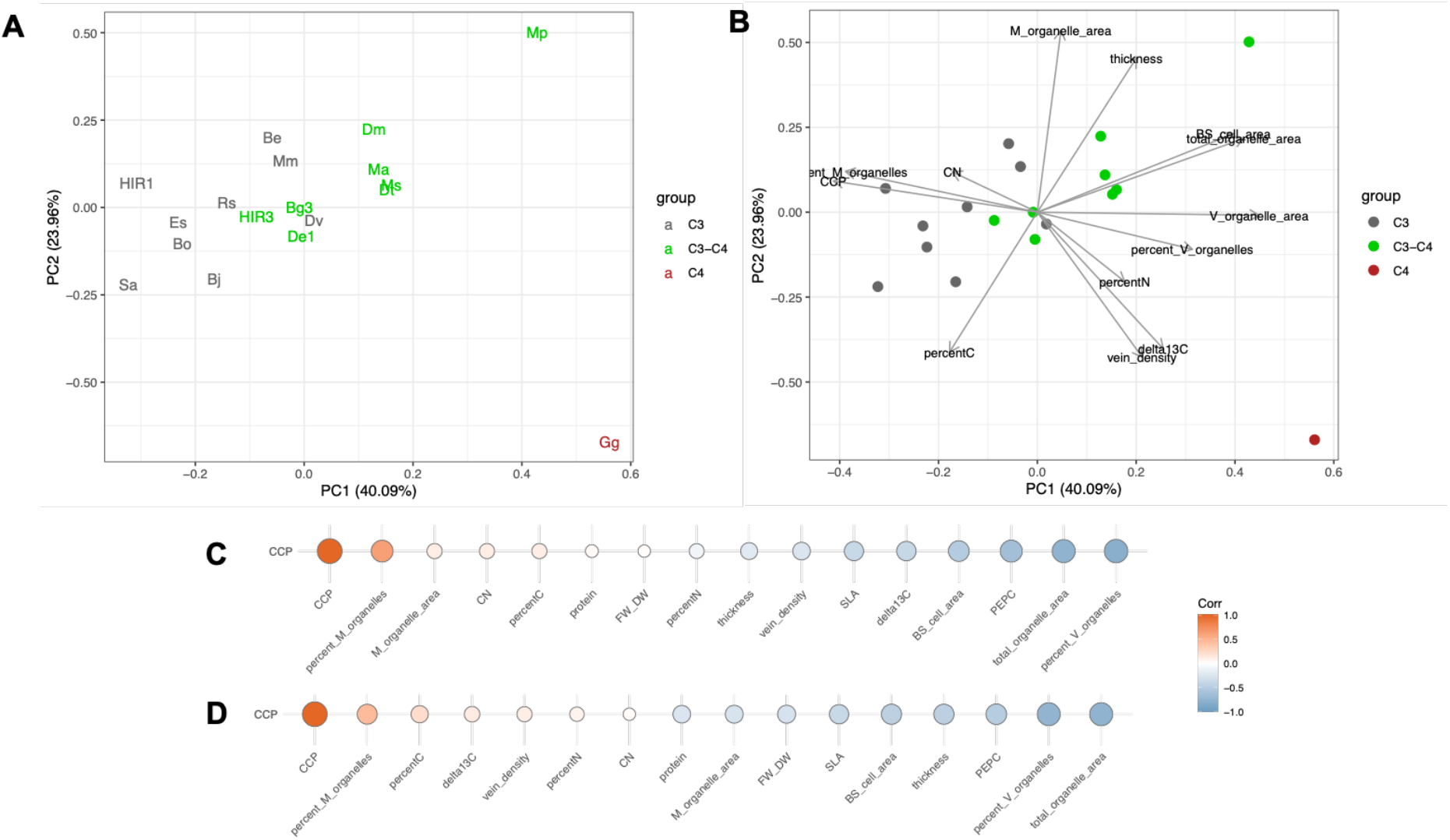
Principal component analysis (PCA) and correlation of the CO_2_ compensation point (CCP) with leaf structural and compositional components. Average values for the selected parameters measured by EA-IRMS, analysis of leaf cross sections by light microscopy. (A) localization of the plant lines in the PCA, (B) PCA including parameter loadings, (C) Pearson correlation coefficients demonstrated as heatmaps using all plant lines, (D) Pearson correlation coefficients demonstrated as heatmaps using only C_3_ and C_3_-C_4_ lines. The tested plant lines were colored according to photosynthesis type as C_3_ (grey), C_3_-C_4_ (green) and C_4_ (red). Species and accessions have been abbreviated for legibility and are provided in Figure 2 and Material and Methods.

Summarising the above-mentioned structural and leaf composition related parameters in a PCA, the C_4_ *G. gynandra* can be separated from the rest of the Brassicaceae plants (Figure 8 A, B). This was mainly driven by high values for D^13^C, vein density and vein orientated organelles in the BS as well as low values for CCP and ICS/M orientated organelles in the BS (Figure 8 A, B). C_3_ and C_3_-C_4_ accessions separated along the same line, a combination of PC1 and PC2, but an overlap between the two groups was nevertheless observed. Correlation of the CCP to the selected components supported the importance of organelle accumulation and orientation in the BS for the activity of the C_4_ as well as the C_3_-C_4_ pathway (Figure 8 C, D; Supplemental figure S2).

## Discussion

Soon after the discovery of C_4_ biochemistry (Hatch and Slack, 1966), *Mollugo verticullata* was described as a species with features intermediate between ancestral C_3_ and C_4_ photosynthesis (Kennedy and Laetsch, 1974). Specifically, this species was characterised as possessing a high number of plastids in the BS in addition to having photorespiratory rates which were between typical values for C_3_ and C_4_ species (Kennedy and Laetsch, 1974). Subsequently, this discovery was followed by an extensive screening experiment which measured the CCP of 439 monocot and 335 dicot species (Krenzer *et al*., 1975). In this experiment, it was found that most species fell within two main groups, of CCPs above 40 ppm (C_3_), and CCPs below 10 ppm (C_4_), respectively. Only two species showed intermediate CCP values between 10 and 40 ppm, the C_3_-C_4_ grass *Panicum milioides* and the C_3_-C_4_ Brassicaceae *Moricandia arvensis*. Following these observations, *M. arvensis* became one of the first model plants for the investigation of the C_3_-C_4_ intermediate photosynthesis type (Bauwe and Apel, 1979; Holaday and Chollet, 1983; Rawsthorne *et al*., 1988*a*,*b*), sometimes also referred to as the *Moricandia* syndrome. Since then, additional species with intermediate photosynthesis features have also been discovered in the Brassicaceae including *M. suffruticosa, M. sinaica, M. spinosa, M. nitens*, and *D. tenuifolia* (Hylton *et al*., 1988; Apel *et al*., 1997). In this study, we describe the physiology, biochemistry and anatomy of 10 species (14 accessions) of C_3_-C_4_ intermediates and 18 C_3_ species (19 accession) from this family.

The results in the current study reveal that carbon concentrating mechanisms have evolved up to five times independently in the Brassiceae tribe (Figures 1, 2), ranging from very efficient C_3_-C_4_ photosynthesis in *D. tenuifolia* and the *Moricandia* genus (*M. suffruticosa, M. arvensis, M. sinaica, M. nitens, M. spinosa*), to relatively weaker CO_2_ concentrating mechanisms in *B. gravinae, D. erucoides* and *H. incana* HIR3 accession. These observations are based on CCP measurements combined with a phylogenetic tree inferred from genomic sequence data across 102 orthogroups. Our study newly identified carbon concentrating mechanisms in an accession of *H. inana* (HIR3) and, for the first time, investigated C_3_-C_4_ features in *D. erucoides* in more detail.

### Brassicaceae display large variation in efficiency of the carbon conservation mechanism

Our survey of CO_2_ concentration mechanisms in the Brassicaceae confirmed that measurements of the CCP represent a valuable tool for the identification of C_3_-C_4_ intermediate plant accessions. However, in contrast to the large screening study by Krenzer et al. (Krenzer *et al*., 1975), we observed gradual changes in the CO_2_ concentrating capacity within the group of potential C_3_-C_4_ intermediates that exhibit CCPs below 40 ppm. Our study therefore supports models claiming that carbon concentrating mechanisms evolved in a smooth manner by additive effects of small fitness gains (Heckmann *et al*., 2013).

The lowest CCPs in the present investigation was measured in *D. tenuifolia* and the C_3_-C_4_ *Moricandia* species. Although various accessions of these species were used in different studies (Hylton *et al*., 1988; Rawsthorne *et al*., 1988*a*; Apel *et al*., 1997; Ueno *et al*., 2003, 2006; Schlüter *et al*., 2017), low CCPs seem to be a ubiquitous trait of these respective species. Moreover, low CCPs in these species were supported by BS specific localisation of the GLDP protein (Hylton *et al*., 1988; Rawsthorne *et al*., 1988*a*; Ueno *et al*., 2003). Further, especially in *D. tenuifolia*, CCP values were observed as very low and close to those typical of C_4_ species. However, PEPC and ^13^C measurements support the fact that C_3_-C_4_ *D. tenuifolia* and *Moricandia* species do not operate a C_4_ cycle.

Four accessions of *B. gravinae* were tested in the present study and all were found to exhibit C_3_-C_4_-like CCP values, indicating stability of this trait in this species. This result is also corroborated by a previous study which has observed GLDP localisation to the BS in this species (Ueno, 2011). Moreover, although variation in the CCP of *D. erucoides* has been previously reported (Apel *et al*., 1997), both accessions of this species characterised in the present study exhibited CCP values typical of C_3_-C_4_ plants.

Interestingly, our analysis included two different accessions of *H. incana*, of which HIR1 clustered with C_3_ species while HIR3 showed a reduced CCP typical of a C_3_-C_4_ intermediate. Variation in the CCP within different populations of the same species had been observed before (Teese, 1995; Apel *et al*., 1997; Lundgren *et al*., 2016; Yorimitsu *et al*., 2019). This could be due to the manifestation of the C_3_-C_4_ phenotype in only some accessions within the species. A recent publication investigating C_3_-C_4_ features in the Chenopodiaceae showed that growth conditions, especially temperature and nitrogen supply, could considerably influence the formation of the glycine shuttle in some species. The CCP was lowest under high temperature and low nitrogen conditions, which was connected to accumulation of the GLDP protein preferentially in the BS (Oono *et al*., 2022). These results in combination with the previous literature support that all species classified as C_3_-C_4_ intermediates in the present study achieve high CO_2_ assimilation efficiency exclusively using the photorespiratory glycine shuttle. Our results also indicate that gradual and even facultative implementation of carbon shuttles between the M and BS are possible and should be considered in future experiments.

In addition to the five independent evolutionary origins of C_3_-C_4_ in the Brassicaceae, *D. muralis* appears to have inherited C_3_-C_4_ intermediacy as it is a tetraploid natural hybrid between the C_3_ species *D. viminea* and the strong glycine shuttle species *D. tenuifolia* (Eschmann-Grupe *et al*., 2003; Ueno *et al*., 2006). In the majority of crossings between C_3_ and C_3_-C_4_ Brassicaceae the physiology of the hybrids was closer to the C_3_ parent (Rawsthorne *et al*., 1998; Ueno *et al*., 2003) and additional evolutionary adjustments might be necessary to further optimise the efficiency of the glycine shuttle (Ueno *et al*., 2003). As such, it has been suggested that hybridisation events could play a major role in the acquisition of the carbon concentrating mechanisms (Kadereit *et al*., 2017). Further, in some grasses, lateral gene transfer has been shown to support the rapid and successful establishment of the C_4_ pathway (Dunning *et al*., 2019*b*). The contribution of such transfer mechanisms for carbon concentrating mechanisms in the Brassicaceae can only be resolved by detailed genome analysis. Such scenarios would nevertheless require donor species that are able to successfully transfer essential features into the receiving genetic background.

### C_3_-C_4_ photosynthesis is associated with reduced Ci and enhanced WUE especially under limiting CO_2_

In the Brassicaceae, the presence of C_3_-C_4_ metabolism did not translate into improved photosynthetic assimilation under ambient environmental conditions (Figure 3). For instance, across the Brassicaceae species analysed in the current study, assimilation rates appeared to be genotype specific rather than related to photosynthesis type under ambient CO_2_. This lack of correlation between assimilation and photosynthesis type has also been previously described in the Chenopodiaceae (Yorimitsu *et al*., 2019).

Interestingly, however, C_3_-C_4_ species in the Brassicaceae adjusted leaf Ci to lower levels compared to C_3_ species in this clade. The difference between these photosynthesis types was marginal under ambient conditions, but became more pronounced under CO_2_ conditions of 200 ppm and lower (Figure 4 C, D). The ability to assimilate CO_2_ at lower Ci translated into higher WUE in the C_3_-C_4_ species compared to the C_3_ species. This increase in WUE observed was underpinned by enhanced assimilation, as stomatal conductance was similar among the C_3_ and C_3_-C_4_ species under all tested conditions (Figures 3 B, D, F). It should however be noted that the differences observed for Ci and WUE between C_3_ and C_3_-C_4_ were small in comparison to the difference between all C_3_ and C_3_-C_4_ species and C_4_ *G. gynandra*, thus underlining the superiority of the C_4_ pathway as a CO_2_ concentrating mechanism compared to C_3_-C_4_ metabolism. Similar observations have been previously observed in *Heliotropium* and *Flaveria*, in which C_3_-C_4_ species achieved WUE values between the C_3_ and C_4_ species with this result also due to higher assimilation rather than modified conductance (Huxman and Monson, 2003; Vogan *et al*., 2007). These results support an advantage of the C_3_-C_4_ pathway in high photorespiratory conditions which cause CO_2_ restriction due to stomatal closure.

Although the C_3_ species *M. moricandioides* appears to be geographically restricted to the Southern Iberian Peninsula, the closely related C_3_-C_4_ species *M. arvensis* has spread into a variety of habitats ranging from Great Britain to Tunisia (Lundgren and Christin, 2017; Perfectti *et al*., 2017); (Supplemental figure S8). Similarly, the C_3_ species *D. viminea* grows mainly in Southern Spain and France, whilst the closely related C_3_-C_4_ species *D. tenuifolia* and *D. muralis* can also be found in colder and wetter more northern locations such as Scandinavia and Scotland (Supplemental figure S8). Thus, in both of these respective lineages, the glycine shuttle appears to be connected to an enlargement of the growth habitat but not necessarily towards specific climatic preferences (Lundgren and Christin, 2017); (Supplemental figure S9). For instance, *D. tenuifolia* often grows as an invasive species occupying sunny, harsh and arid sites (Nicoletti *et al*., 2007) in which water, nutrient and temperature conditions can change rapidly. In competitive habitats, C_4_ species have been shown to profit especially from their ability for fast biomass accumulations under favourable conditions (Christin *et al*., 2014; Knapp *et al*., 2020), and efficient conservation of photorespiratory CO_2_ possibly contributed to fast growth and establishment of C_3_-C_4_ species in marginal habitats.

Ecological studies which have investigated the adaptation of C_3_-C_4_ species to specific environmental conditions are unfortunately still rare (Oono *et al*., 2022). However, the present results show that C_3_-C_4_ metabolism evolved in species from different habitats and in different genetic backgrounds in the Brassiceae group and thus indicate benefits of this trait under a wide range of conditions. Ecological preferences of the C_3_-C_4_ species are still dependent on their C_3_ ancestors (Lundgren and Christin, 2017), but installation of the glycine shuttle seems to have broadened their niches. Furthermore, some C_3_-C_4_ species show high plasticity in leaf morphological traits. *M. arvensis* leaves had lower CCPs and higher WUE under hotter and more arid summer conditions than in milder spring climate (Gomez et al., 2020). A similar phenomenon was recently described for *Chenopodium album* where selected accessions developed low CCPs under high temperature and low nitrogen conditions (Oono *et al*., 2022). C_3_-C_4_ species could generally be associated with harsh environments experiencing large ranges of temperature and water availability, thereby possibly profiting from high environmental flexibility of the trait.

Knowledge about the distribution of species with glycine shuttle metabolism is generally still limited to studies among relatives of C_4_ species. As such, very few larger surveys have been performed which assess CCP across a broad range of plant species (Krenzer *et al*., 1975; Apel *et al*., 1997). This is mainly due to the dependence on gas exchange equipment and time-consuming measurements. It is therefore assumed that the frequency of species with weaker carbon concentrating mechanisms is strongly underestimated (Sage *et al*., 2011*b*; Lundgren, 2020). Identification of CCPs characteristic to C_3_-C_4_ species in a *H. incana* accession and recently in some *C. album* accessions (Yorimitsu *et al*., 2019) support this hypothesis. As such, faster methods for identification of C_3_-C_4_ intermediates could help to close this knowledge gap. Here, our correlation analysis showed that measurements of assimilation at low CO_2_ are sufficient for detection of C_3_-C_4_ phenotype and would save considerable time as appose to having to calculate CCP by measuring assimilation across a range of CO_2_ concentrations (Figure 4 D). For example, a very strong positive correlation in the present results was found to exist between CCP and assimilation rate at 50 ppm CO_2_ which is close to the CCP of C_3_ species. High and significant negative correlation to CCP also existed for WUE under CO_2_ conditions of 200 ppm and below. As in our experiments assimilation generally correlated positively with photosynthesis efficiency Fv’/Fm, fluorescence combined with stomatal conductance measurements could possibly also be used in a fast initial screening experiments for identification of C_3_-C_4_ intermediates in the future.

### Reduction in CCP correlates negatively with organelle accumulation and arrangement in BS

Although a gradual change in various physiological parameters was observed during transition between C_3_ and C_3_-C_4_ photosynthesis, more distinct changes were found in BS structural data. For instance, all plants from the C_3_-C_4_ group possessed enhanced BS area occupied by organelles in the centripetal position and a higher total organelle area per BS cell compared to C_3_ species (Figures 7, 8). This underlines the importance of anatomical features for carbon recapturing mechanisms. A strong correlation between reduction in CCP and increased organelle accumulation facing the vein in the BS was also previously observed in interspecific hybrids between *D. tenuifolia* (C_3_-C_4_) and *R. sativus* (C_3_) (Ueno *et al*., 2003). The BS structural features appeared to be genetically encoded and is inherited independently from the GLDP localisation (Ueno *et al*., 2003). Residual expression of the *GLDP* was also observed in M cells of C_3_-C_4_ intermediate *Flaveria* species (Schulze *et al*., 2013). This shows that structural modifications can underpin an effective CCP without complete suppression of GLDP in the M cells.

C_3_-C_4_ intermediates in our study contained several layers of M cells such that many do not directly border BS cells. This would mean that complete absence of GDC activity in the M cells would require transport of photorespiratory glycine through other M cell layers prior to entering the BS for metabolisation. However, accumulation of mitochondria in the BS might create a glycine sink supporting glycine diffusion to the BS and partial reduction of M GLDP expression would enforce this shuttle. TEM studies of centripetally localised organelles from C_3_-C_4_ Brassiceae (Ueno *et al*., 2006; Schlüter *et al*., 2017), Asteraceae (McKown and Dengler, 2007), Boraginaceae (Muhaidat *et al*., 2011), Scrophulariaceae (Khoshravesh *et al*., 2012), Arthropogoninae (Khoshravesh *et al*., 2016) and Chenopodiaceae (Yorimitsu *et al*., 2019) have shown a close arrangement of mitochondria and chloroplasts. Thus, BS ultrastructure seems to play a major role in prevention of photorespiratory CO_2_ and NH_3_ loss and in improvement of leaf carbon and possibly also nitrogen economy.

In contrast to the C_4_ specie in our study, BS cells in C_3_-C_4_ Brassicaceae species exhibited organelles facing the ICS and M cells (Figure 7, Supplemental figure S7). This amount of ICS/M cell facing organelles decreased in C_3_-C_4_ species with higher carbon concentrating efficiency. Our results suggest that accumulation of centripetal organelles and reduction of peripheral organelles are not necessarily regulated by the same process. Additional structural features of C_4_ species such as enlarged BS cell area and higher vein density did not differ between the tested C_3_ and C_3_-C_4_ Brassicaceae species. Further, leaf thickness, SLA and FW/DW ratios were also not different between the leaves of the C_3_, C_3_-C_4_ and C_4_ species (Figure 7, Supplemental figure S3). Thus, despite leaf anatomy and BS architecture being important requirements for evolution of carbon concentrating mechanisms (Christin *et al*., 2013), modifications in leaf succulence parameters do not appear to be essential for efficient photorespiratory carbon concentration in the Brassicaceae. Plasticity in morphological parameters could also play a role in further evolution towards the C_4_ leaf and thus be connected to the absence of C_4_ photosynthesis in this plant group (Schlüter *et al*., 2017).

### Brassicaceae C_3_-C_4_ metabolism had only minor influence on leaf steady-state metabolite patterns

Organelle accumulation in the BS and shift of GDC activity to this tissue influences leaf biochemistry (Rawsthorne, 1992; Schlüter *et al*., 2017). For instance, a lack or reduction of GDC activity in the M causes accumulation of photorespiratory glycine and its transport along the concentration gradient to the BS where it is efficiently recaptured in the numerous BS chloroplasts by rubisco. Similar mechanisms have evolved independently in different phylogenetic backgrounds and uneven distribution of the GLDP protein has been confirmed for numerous plant species (Schlüter and Weber, 2020). The GDC reaction, however, also utilizes NADH and releases NH_3_ alongside CO_2_, further metabolic balancing between the two cell types. Nevertheless, beyond GLDP localisation, not much is known about the cell specific metabolism or the nature of additional metabolite shuttles in C_3_-C_4_ Brassicaceae.

If metabolite exchange between M and BS cells is realised by a concentration gradient, high concentrations of these transported metabolite would be expected in the leaves (Leegood and Von Caemmerer, 1989). Although, it should be noted that high metabolic flux and cell-specific metabolite accumulation might mask these gradients in total leaf extracts. Our metabolite analysis did not identify preferential metabolite shuttles operating across all C_3_-C_4_ Brassicaceae species. Steady-state glycine concentrations were generally enhanced in the C_3_-C_4_ species compared to C_3_ species, supporting the hypothesis that glycine is transported from the M cells to the BS for decarboxylation. High glycine was, however, also found in leaves of the C_3_ *D. tenuisiliqua* and the C_4_ species *G. gynandra* indicating that glycine accumulation is not a distinct C_3_-C_4_ features (Figure 6A). Further uncertainty exists around the metabolites transported back from BS to M for rebalancing of carbon, nitrogen and energy metabolism (Borghi *et al*., 2022). Beside glycine, serine accumulation also exhibited a negative correlation with CCP values (Figure 5 D). This strongly supports the involvement of serine as a metabolite transported back from the BS to the M cells (Rawsthorne, 1992; Mallmann *et al*., 2014), although variation in serine levels suggest that the contribution of serine transport could vary between the different accessions.

Strong variation between the individual accessions also existed for other shuttle metabolite candidates. Modelling approaches have previously predicted the involvement of glutamate, α-ketoglutarate, α-alanine, pyruvate, aspartate and malate in shuttling processes for rebalancing of nitrogen metabolism between M and BS (Mallmann *et al*., 2014). Malate and aspartate could also be involved in rebalancing of reducing power between the two cell types (Johnson *et al*., 2021). Contribution of glutamine/glutamate and asparagine/aspartate to intercellular shuttles were suggested for the C_3_-C_4_ species *Flaveria anomala* (Borghi *et al*., 2022). Here, enhanced levels of these various metabolites could be observed in some, but not all C_3_-C_4_ accessions (Figure 6). For example, high concentrations of malate, aspartate and glutamate were found in species displaying very low CCPs such as *M. arvensis* and *D. tenuifolia*. Interestingly, the C_3_-C_4_ *Moricandia* species which supposedly share a single C_3_-C_4_ evolutionary origin also showed strong variation in the metabolite pattern. A similar absence of main shuttle metabolites has also been described for C_3_-C_4_ *Flaveria* species (Borghi et al., 2022). Our data generally support the hypothesis that multiple metabolites are transported between M and BS (Schlüter *et al*., 2017; Borghi *et al*., 2022). The contribution of the different metabolites could differ in the individual accessions depending on genetic as well as environmental influences. Such a multitude of solutions indicates that metabolite and energy balancing does not represent a limiting step during evolution of carbon concentrating pathways.

To date, enzyme localisation studies have mostly focussed on the GLDP protein, and much less is known about whether other reactions are shifted to the BS in C_3_-C_4_ species. In *M. arvensis*, other tested photorespiratory enzymes such as glycolate oxidase, serine hydroxymethyl transferase and other subunits of the GDC complex were present in both cell types (Morgan *et al*., 1993). Enzyme activities in *M. arvensis* in M and BS enriched fractions were also equally distributed for glyoxylate aminotransferases, glycolate oxidase and hydroxypyruvate reductase (Rawsthorne *et al*., 1988*b*) supporting the crucial role of uneven distribution for glycine shuttle operation in *M. arvensis*. On the other hand, all GDC subunits were preferentially expressed in the BS in C_3_-C_4_ *Flaveria* and *Panicum* species (Morgan *et al*., 1993). Shifting of additional photorespiratory steps could considerably influence the metabolite shuttles. In our study, some accessions, especially *D. erucoides* and *D. tenuifolia*, showed high levels of glycolate and glycerate. Interestingly, intercellular transport of glycerate and glycolate was predicted in a constraint-based modelling approach for weak carbon concentrating mechanisms on the evolutionary path to C_4_ photosynthesis (Blätke and Bräutigam, 2019). Exchange of these metabolites between M and BS would reduce the need for intercellular nitrogen recycling (Borghi *et al*., 2022). Part of the photorespiratory metabolites could also feed into additional pathways in the BS. It has been estimated that 1-5 % of the photorespiratory glycine and about 30% serine can be metabolised outside the photorespiratory cycle in processes such as protein biosynthesis (Busch *et al*., 2018). The high organelle accumulation would increase the demand for protein synthesis in the C_3_-C_4_ BS. Furthermore, the BS is also responsible for loading of assimilation products into the phloem, and part of the carbon and nitrogen transported into the BS by the glycine shuttle could support metabolite export to the sink tissue of the plants.

In our study, none of the tested C_3_-C_4_ species showed significantly increased PEPC activity or reduced ^13^C levels when compared to the C_3_ group (Figure 7 C). Similarly, it has been previously reported that there is no evidence for a C_4_ cycle operating beside the glycine shuttle in *M. arvensis* (Holaday and Chollet, 1983; Hunt *et al*., 1987). Thus, despite the multiple origins of C_3_-C_4_ photosynthesis in the Brassicaceae, no species appears to have evolved any substantial C_4_ biochemistry.

## Conclusions

Our survey of the Brassicaceae family revealed gradual differences in the carbon concentrating mechanisms which reached very low CCPs of around 12 ppm in *D. tenuifolia*. Here, reduction in CCP was generally associated with organelle arrangement in the BS. Thus, elucidation of regulatory mechanisms underlying organelle multiplication and arrangement in the BS appear to be crucial for engineering an efficient glycine shuttle pathway into the leaf.

The situation in the Brassicaceae appears to be unique considering the fact that in this clade the glycine shuttle has evolved multiple times despite no evidence of C_4_ photosynthesis having evolved in this plant family. All C_3_-C_4_ classified species belong to the Brassiceae tribe appear to have lost one GLDP gene copy suggesting that this gene loss event has facilitated evolution of the glycine shuttle (Schlüter *et al*., 2017). Metabolite data indicated that glycine and serine are involved in metabolite shuttling, but that additional individual shuttles could be active in the different C_3_-C_4_ plant accessions. Our results suggest that multiple solutions for intercellular metabolite balancing are possible. In the investigated accessions, carbon concentration by recapturing of photorespiratory CO_2_ seems to be a stable and independent fitness trait distinct from C_4_ photosynthesis in physiology, biochemistry and anatomy.

Carbon concentrating mechanisms have mainly been associated with hot and dry conditions that promote photorespiration. However, the geographical distribution of the C_3_-C_4_ Brassiceae species shows that this phenotype can enable tolerance of a broad range of habitats, especially sites with fast changing temperature, water and nutrient conditions. Such metabolic plasticity could also be advantageous in crop species challenged by climate changes. Crops like *B. napus* or *B. oleaceae* are closely related to described C_3_-C_4_ species and would be prime targets for transfer of this trait. Recent progress in sequencing the genomes of these species and related species in the Brassicaceae (Guerreiro *et al*., 2023) can help to identify the molecular mechanisms behind BS specific C_3_-C_4_ architecture and biochemistry.

## Supplementary material

**Supplemental table S1**: Origin of the seed material

**Supplemental table S2:** Physiological data measured with the Li-COR 6800 (average per accession, standard deviation, HSD group)

**Supplemental table S3**: Metabolite data from GCMS measurements (average per accession, standard deviation, HSD group)

**Supplemental table S4**: Data from light microscopy; EA-IRMS, and PEPC enzyme assay (average per accession, standard deviation, HSD group)

**Supplemental figure S1**: Heatmap Pearson correlation matrix for physiological gas exchange and fluorescence data

**Supplemental figure S2**: Heatmap Pearson correlation matrix for CO_2_ compensation points and metabolite data from GC-MS analysis

**Supplemental figure S3**: Heatmap Pearson correlation matrix for CO_2_ compensation point, structural parameters from light microscopy analysis, leaf composition data from EA-IRMS analysis and PEPC enzyme activity

**Supplemental figure S4**: Box Whisker plot for gas exchange and fluorescence data summarised per photosynthesis type

**Supplemental figure S5**: Box Whisker plot for metabolite data summarised per photosynthesis type

**Supplemental figure S6**: Box Whisker plot for structural and leaf composition data summarised per photosynthesis type

**Supplemental figure S7**: Micrographs of bundle sheath cross sections

**Supplemental figure S8**: Geographical distribution maps of selected Brassicaceae species in Europe/Northern Africa

**Supplemental figure S9**: Climate data associated with habitats of selected Brassicaceae species

## Acknowledgment

The authors would like to thank the gardener team for looking after our plants in the HHU greenhouses. The technical assistance of Katrin Weber, Maria Graf, Elisabeth Klemp and Dominik Brilhaus in the metabolite analysis was also greatly appreciated. We are grateful to Stephanie Krey for assistance with DNA extraction and Perlina Lim for help during the setup of the microscopy experiments. Sebastian Triesch is thanked for his helpful comments on the manuscript.

## Authors contributions

APMW, BS and US initiated and planned the experiments. US, JWB, MM and CK contributed data on plant physiology, biochemistry and structure. PW measured and analysed the plant metabolites. RG and BS provided sequence data and the phylogenetic tree. US wrote the manuscript. JWB, MM, RG, BS, PW and APMW edited the manuscript.

